# Automated assembly of molecular mechanisms at scale from text mining and curated databases

**DOI:** 10.1101/2022.08.30.505688

**Authors:** John A. Bachman, Benjamin M. Gyori, Peter K. Sorger

## Abstract

The analysis of ‘omic data depends heavily on machine-readable information about protein interactions, modifications, and activities. Key resources include protein interaction networks, databases of post-translational modifications, and curated models of gene and protein function. Software systems that read primary literature can potentially extend and update such resources while reducing the burden on human curators, but machine-reading software systems have a high error rate. Here we describe an approach to precisely assemble molecular mechanisms at scale using natural language processing systems and the Integrated Network and Dynamical Reasoning Assembler (INDRA). INDRA identifies overlaps and redundancies in information extracted from published papers and pathway databases and uses probability models to reduce machine reading errors. INDRA enables the automated creation of high-quality, non-redundant corpora for use in data analysis and causal modeling. We demonstrate the use of INDRA in extending protein-protein interaction databases and explaining co-dependencies in the Cancer Dependency Map.

## INTRODUCTION

Molecular biology is characterized by a sustained effort to acquire and organize mechanistic information about the molecules governing the behavior of cells, tissues and organisms (Craver and Darden, 2013). “Mechanism” is used rather loosely in this context, since it operates on multiple scales from the structural transitions of individual molecules to the myriad interactions mediating signal transduction, but it is generally understood to involve a description of the properties, modifications and behaviors of biomolecules in terms of physical and chemical principles. Individual mechanistic discoveries are reported in the biomedical literature, which, with over 3 x 10^7^ articles indexed in PubMed as of 2022, constitutes a substantial public investment and an essential source of knowledge. However, results in research papers are generally described in natural language designed for human – not machine – consumption. As the literature has grown, and methods of experimental data collection become more diverse, it has become impossible for any individual scientist to acquire all of the background knowledge necessary to be an expert in a particular problem and fully interpret experimental results (Forscher, 1963). Biomedicine is therefore faced with a substantial problem of knowledge aggregation, harmonization, and assembly.

The bioinformatics community has actively worked to make knowledge more accessible by curating information about molecular mechanisms in a machine readable form suitable for computational data analysis (Ashburner et al., 2000; Fabregat et al., 2018; Perfetto et al., 2016; Schaefer et al., 2009). This has led to the creation of standard representation languages (Demir et al., 2010; Hucka et al., 2003), and databases that aggregate curated knowledge from multiple primary sources (Cerami et al., 2011; Jensen et al., 2009; Türei et al., 2016). Curated databases form the backbone of many widely used methods of high-throughput data analysis, including gene set and pathway enrichment, and prior knowledge-guided network inference (Babur et al., 2021; Dugourd et al., 2021). However, creation of these database has largely involved human curation of the literature, which is costly and difficult to sustain (Bourne et al., 2015). As a result, most databases and online resources are incomplete; for example, the creators of Pathway Commons (which aggregates pathway knowledge from 22 primary human-curated databases) have estimated that their resource covers only 1-3% of the available literature (Valenzuela-Escárcega et al., 2018). At the same time, databases such as Pathway Commons contain redundant or conflicting information about the same sets of mechanism because assembling knowledge into a coherent whole remains difficult and is currently dependent on additional human curation. Compounding these difficulties is the increasing volume of published scientific articles and the fact that curation standards and languages evolve along with methods of data collection and analysis, making on-going maintenance of a previously curated resources necessary to prevent them from becoming obsolete.

Automated extraction of mechanistic information through literature mining has the potential to address many of the challenges associated with manual curation (Ananiadou et al., 2015). However, the precision of machine reading systems is still lower than that of human curators, particularly for complex relationships that underly many statements about mechanism (Allen et al., 2015; Islamaj Doğan et al., 2019; Madan et al., 2019). Nevertheless, at the current state of the art, machine reading can extract simple relations (e.g., post-translational modifications and binding and regulatory events) at literature scale and with reasonable reliability. A variety of text mining systems have been developed, each with different designs, strengths, and weaknesses, but common steps include grammatical parsing of sentences, named entity recognition and normalization, also called grounding (i.e., associating entities with a standardized identifier in controlled vocabularies such as HGNC), and event extraction (identifying interactions, transformations or regulations among grounded entities). Much of the research to date in text mining for biology has focused on small-scale studies for method validation, but a handful of efforts have aimed to create large-scale resources available for use in data analysis by the broader computational biology community (Van Landeghem et al., 2013; Yuryev et al., 2006). We speculate that the reliability of machine reading could be increased by combining the results of *multiple* distinct systems in a principled manner, but few such combined approaches have been described thus far.

What is still needed are computational tools for the large-scale assembly of both text-mined *and* curated mechanisms in databases to generate knowledge resources with mechanistic detail and genome scale. Human-generated resources such as Reactome (Fabregat et al., 2018) aspire to this, but would benefit in scope and currency from human-in-the-loop collaboration with machines. However, machine assembly must overcome not only errors in grounding and event extraction but also the challenges associated with combining noisy information about mechanisms at different levels of specificity. Users of this information may have different end goals, but have a common need for reliable networks and models that can be used to investigate mechanisms at the level of the individual reactions, mutations, or drug-binding events—something currently possible on a smaller scale using dynamical systems analysis (Lopez et al., 2013) and logic-based modeling (Saez-Rodriguez et al., 2009). These more mechanistic networks and models contrast with existing genome-scale networks that commonly involve unsigned node-edge graphs that aggregate diverse types of interactions (genetic, physical, co-localization, etc.) using the simplest possible abstraction.

We previously described a software system, the Integrated Network and Dynamical Reasoning Assembler (INDRA) that automates the use of curated natural language text to create computational models that can be executed using dynamical, logic-based, or causal formalisms (Gyori et al., 2017). For example, we have previously used INDRA to convert “word models” expressed in simplified declarative text (e.g., “*Active ATM activates p53*. *Active p53 transcribes MDM2 etc*.”) into dynamical ODE-based models. A key feature of INDRA is that it uses an intermediate representation to decouple the process of knowledge collection from the construction of specific models (**Figure 1A**). More specifically, INDRA normalizes mechanistic information expressed in natural (English) language into a high-level intermediate machine representation called Statements. Statements can then be used directly to create executable models, for example in rule-based languages such as BioNetGen or PySB. The current taxonomy of INDRA Statements accounts for the types of biomolecular processes most commonly involved in intracellular biological networks and signal transduction (e.g., post-translational modifications, positive and negative regulation, binding, transcriptional regulation, etc.) but is extensible to other domains of natural science.

**Figure 1.**
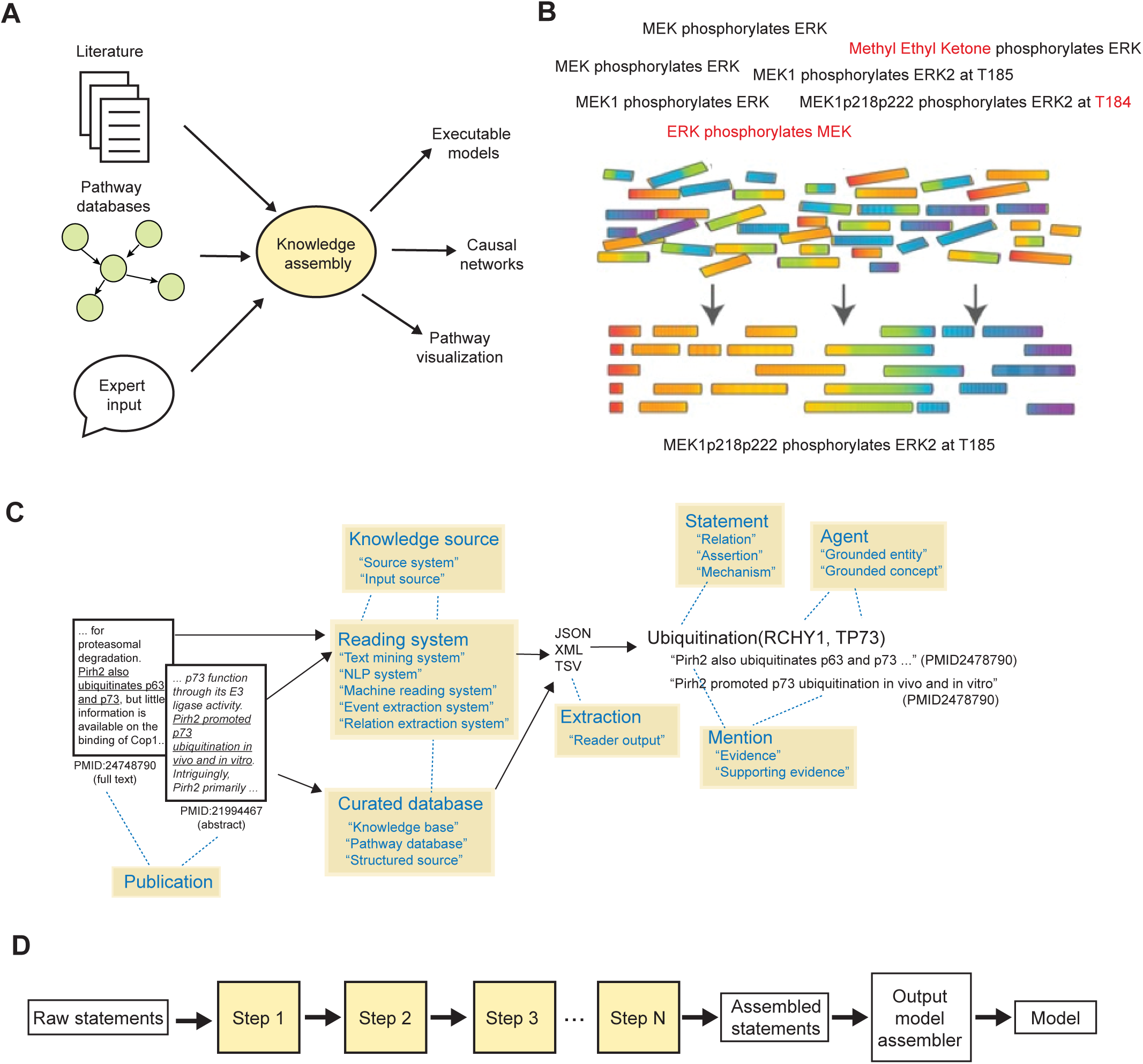
Conceptual overview of knowledge assembly. (A) Assembly of models from diverse knowledge sources. Structured (pathway databases) and unstructured (literature, expert input in natural language) biological knowledge is converted into machine-readable, mechanistic fragments. These fragments must be assembled into a coherent corpus before generation of specific models for data analysis. (B) Mechanistic “fragments” capture incomplete but overlapping aspects of an underlying molecular mechanism (here, the phosphorylation of ERK by MEK). Fragments may also contain errors (highlighted in red). Assembly involves identifying relationships between fragments in order to arrive at a consensus representation that captures available information. (C) Artifacts involved in the collection of mechanisms from knowledge sources by INDRA, and their representation as INDRA Statements. Yellow boxes show key terminology used to refer to different artifacts with additional synonyms provided in quotes. (D) INDRA knowledge assembly transforms raw statements into assembled statements from which models can be generated. The individual steps of the assembly pipeline (Steps 1 to N, yellow background) operate on INDRA Statements and are configurable from a library of built-in or user-defined functions.

Here we describe a greatly expanded set of computational tools and use cases, implemented within the INDRA architecture, for combining mechanistic information obtained at scale from primary research publications. This is a substantially more challenging task than the conversion of short, controlled, declarative text into ODE models that we described previously (Gyori et al., 2017). We accomplished reading at scale by combining the results of multiple reading systems with curated mechanisms from a wide range of databases and structured knowledge sources. Used in this way, INDRA identifies duplicate and partially overlapping Statements, allowing for automated assembly of mechanistic fragments into a nonredundant and coherent set of interactions and subsequently into large-scale knowledge assemblies for use in biocuration and data analysis. We illustrate the end-to-end assembly procedure with INDRA by processing publications specifically relevant to human genes and integrate this information with publicly available databases to create a corpus of ∼ 900,000 unique and specified interactions among human proteins. We found that overlap between different machine reading systems was surprisingly small (highlighting both the readers’ complementarity and their limitations), but for a given Statement, the existence of supportive evidence from multiple systems was informative of reliability. We used manual curation to analyze the error and overlap characteristics of multiple machine reading systems and, using this data, we developed predictive models that estimate the technical reliability of text-mined extractions in the form of a “belief score”. To evaluate the utility of machine-extracted mechanisms we used the INDRA-assembled corpus of Statements to prioritize protein-protein interactions for curation that are not yet captured in the widely used structured knowledgebase, BioGRID (Oughtred et al., 2019). Finally, we used the same assembled corpus to identify and explain gene dependency relationships in the Cancer Dependency Map (DepMap) dataset (Meyers et al., 2017; Tsherniak et al., 2017). In this case, an INDRA-assembled network served helped determine statistically significant codependencies between genes, thus allowing for the detection of new codependencies in the context of cancer. INDRA also provided possible mechanistic explanations rooted in the scientific literature for observed DepMap codependencies.

## RESULTS

Automated assembly of large knowledge bases from curated databases and machine reading systems raises a series of interconnected issues not arising in the conversion of curated natural language text to machine readable mechanisms (**Figure 1A**) (Gyori et al., 2017). In particular, each source of information yields many mechanistic fragments that capture only a subset of the underlying process, often at different levels of abstraction. For example, one source might describe the MEK1 (HUGO name *MAP2K1*) phosphorylation of ERK2 (*MAPK1*) on a specific threonine residue (T185), whereas another source might describe the same process at the protein family level, stating that MEK phosphorylates ERK, without mentioning a specific isoform, residue or site position (**Figure 1B**). Individual mechanisms obtained from machine reading are not only fragmented, they also include different types of technical errors that must be overcome (**Figure 1B**, red font). One analogy for assembling pathways from mechanistic fragments is the assembly of a full genome sequence from many noisy, overlapping sequencing reads (**Figure 1B**). The goal of knowledge assembly is similarly to achieve a best “consensus” representation of the underlying processes, incorporating as much mechanistic detail as possible while minimizing errors. Ultimately, the process is expected to yield computational approaches for finding truly missing or discrepant information, by analogy with variant calling.

### Box 1. Representing knowledge captured from multiple sources in INDRA (**Figure 1C**)

Scientific *publications* contain descriptions of mechanisms (interaction, regulation, etc.) among biological entities. These descriptions can be extracted either by human experts and captured in *curated databases* or extracted automatically by *reading systems* using natural language processing. Collectively, reading systems and curated databases serve as *knowledge sources* for INDRA. These *extractions* are made available by knowledge sources in a variety of custom machine-readable formats such as JSON, XML and TSV. INDRA processes such extractions into INDRA Statements. Each *Statement* represents a type of mechanism (e.g., Ubiquitination), and has multiple elements, including *Agents* representing biological entities such as proteins or small molecules, and potentially other mechanistic detail such as an amino acid residue for a modification. Each *Statement* can be supported by one or more *mentions,* each representing a single curated database entry or a single extraction by a reading system from a sentence in a given publication. Mentions are represented by INDRA as Evidence objects that have a multitude of properties representing rich provenance for each mention, including the source sentence and the identifiers of the source publication.

Our preliminary studies identified multiple technical and conceptual problems that needed to be addressed to assemble coherent knowledge at scale. These included (i) inconsistent use of identifiers for biological entities among different sources, (ii) full or partial redundancy between mechanisms, and (iii) technical errors in named entity recognition and relation extraction. Such problems are particularly salient when integrating literature-mined interactions, but they also exist when aggregating interactions from multiple curated databases, due to differences in curation practices. For example, in Pathway Commons v12 there are at least eight different curated representations of the process by which *MAP2K1* phosphorylates *MAPK1*, each at a different level of detail (**Figure S1A).** We developed a set of INDRA algorithms for addressing each of these assembly challenges. These algorithms are general-purpose and can be configured to support a wide range of modeling applications (**Figure 1D**), as illustrated in the following examples of machine reading, assembly, and data analysis.

### INDRA integrates mechanisms from pathway databases and machine reading

We used six machine reading systems, Reach (Valenzuela-Escárcega et al., 2018), Sparser (McDonald et al., 2016), MedScan (Novichkova et al., 2003), TRIPS/DRUM (Allen et al., 2015), RLIMS-P (Torii et al., 2015), and the ISI/AMR system (Garg et al., 2016) to process 567,507 articles (using full-text content when available, and allowed by copyright restrictions, and abstracts otherwise; **Table 1**) curated as having relevance to human protein function (see Methods). Reader output was normalized to INDRA Statements, yielding ∼5.9·10^6^ unassembled or “raw” Statements (**Figure 2A).** These were combined with approximately 7.3·10^5^ INDRA Statements extracted from structured sources such as Pathway Commons and the BEL Large Corpus; this used the previously described extraction logic (Gyori et al., 2017) but extended to multiple additional sources including SIGNOR (Perfetto et al., 2016). In combination, reading and databases yielded a total of ∼6.7·10^6^ raw Statements (the end-to-end assembly is illustrated schematically in **Figure 2A)**. In what follows, we refer to the resulting set of assembled INDRA Statements as the *INDRA Benchmark Corpus*.

**Figure 2.**
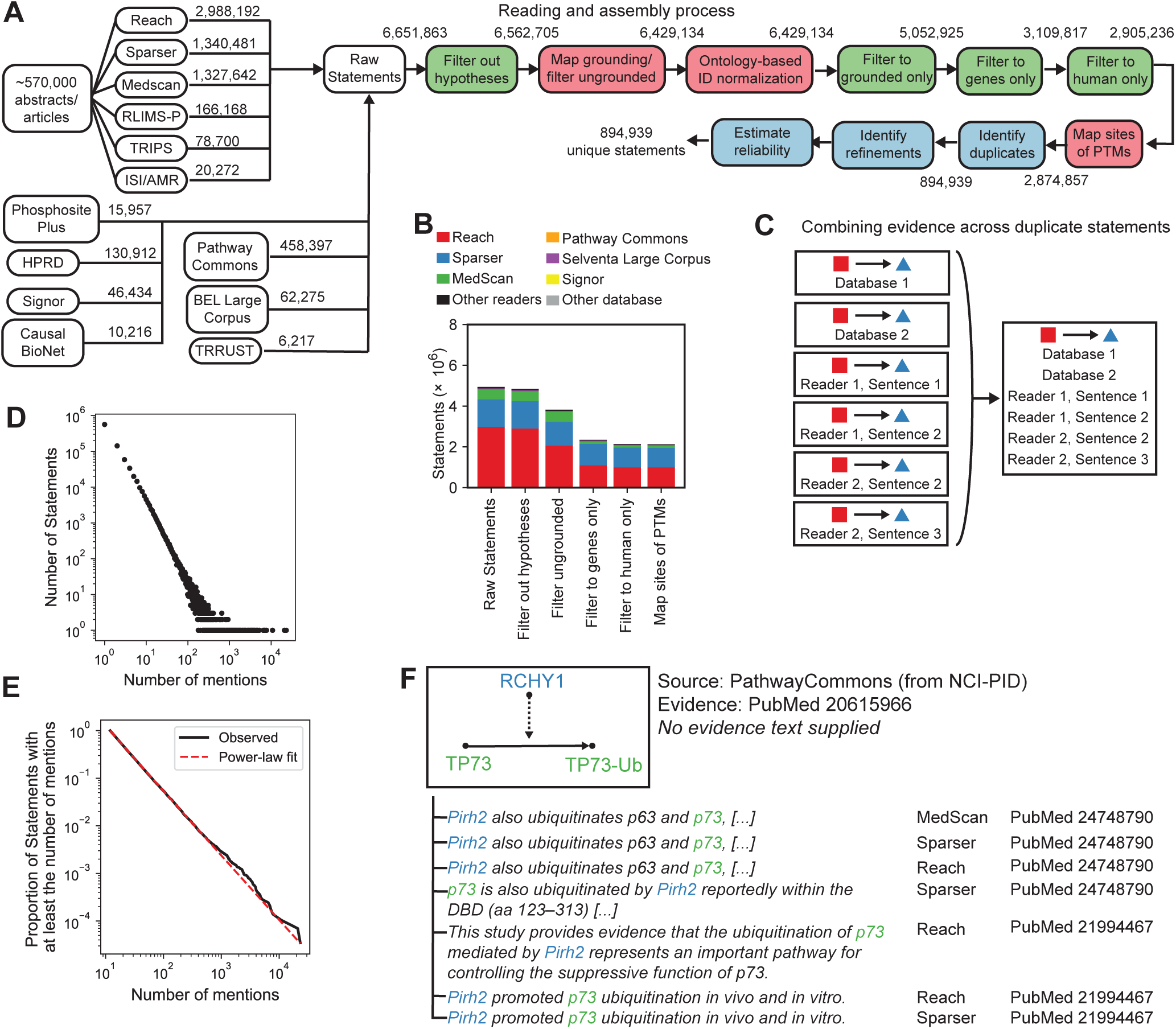
The INDRA knowledge assembly pipeline used to create a Benchmark Corpus. (A) The INDRA assembly pipeline for the Benchmark Corpus. The pipeline starts with ∼570 thousand publications processed by multiple reading systems, as well as structured database sources including Pathway Commons and SIGNOR. Raw Statements extracted from these sources proceed through filtering (green), error-correction (red), and assembly (blue) steps. (B) Number of INDRA Statements, by source, at key stages of the assembly pipeline shown in panel (A). (C) Combining duplicate Statements. INDRA identifies raw Statements that are identical and creates a single unique Statement with all of the associated mentions. (D) Distribution of mention counts (including both mentions in text and database entries) across all Statements in the Benchmark Corpus. Each point in the scatterplot represents the number of Statements with a given number of mentions. (E) Complement cumulative distribution of Statements as a function of the number of mentions supporting them (black) and the maximum likelihood estimate of a power-law fit to the distribution (red). (F) Assembly of Statements enriches curated mechanisms in pathway databases with literature evidence from text mining. Here, a reaction in Pathway Commons represents the ubiquitination of TP73 (p73) by the ubiquitin ligase RCHY1 (Pirh2). Reach, Sparser and MedScan each extract statements matching the one from Pathway Commons and provide references to PubMed identifiers and specific evidence sentences as provenance.

**Table 1:**
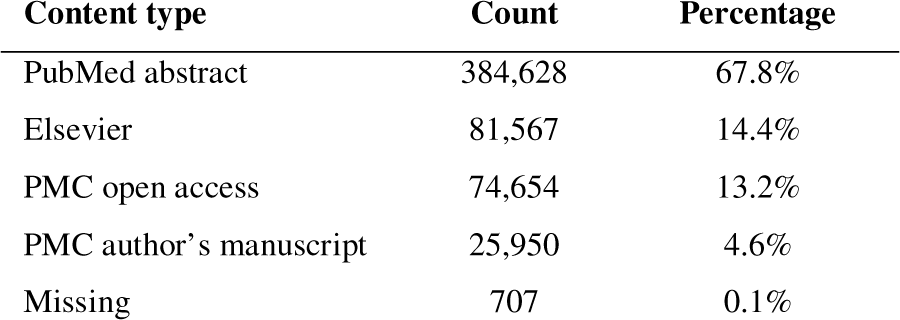
Distribution of content types for literature corpus

| Content type | Count | Percentage |
| --- | --- | --- |
| PubMed abstract | 384,628 | 67.8% |
| Elsevier | 81,567 | 14.4% |
| PMC open access | 74,654 | 13.2% |
| PMC author’s manuscript | 25,950 | 4.6% |
| Missing | 707 | 0.1% |

After collecting information from each source, a series of normalization and filtering procedures were applied (green and red boxes, **Figure 2A**). These processing steps have been combined into a custom computational pipeline but are also available as individual and reusable software modules in INDRA. First, we removed Statements that were supported by mentions indicative of a hypothesis rather than an assertion (for instance, including sentences phrased as “we tested whether…”). Next, “grounding mapping” was performed to correct systematic errors in named entity normalization, which often arise due to the ambiguity of biomedical naming conventions. INDRA integrates both a manually-curated mapping table to fix entities frequently mis-identified by reading systems (described in detail in (Bachman et al., 2018)), and a set of machine learned models to perform disambiguation based on text context (by integrating the Adeft (Steppi et al., 2020) and Gilda (Gyori et al., 2022) systems). “ER” is an example of a common but ambiguous entity: it can stand for endoplasmic reticulum, estrogen receptor, estradiol receptor, emergency room, and a variety of other entities and concepts depending on context. As currently implemented, Reach, Sparser and other reading systems ground “ER” deterministically to a single identifier (e.g., estrogen receptor) irrespective of context. In contrast, the machine learned disambiguation models integrated into INDRA predict the most likely meaning of entities such as ER based on surrounding text; this is then used to current the results of text reading systems.

The next step of the grounding mapping process standardizes identifiers for individual entities using a network of cross-references between equivalent identifiers in different namespaces (Figure S2A). This addresses the opposite problem from the one described above (i.e., one name corresponding to multiple entities), namely that a single entity can have multiple identifiers in different namespaces, and these identifiers can be assigned inconsistently across machine reading systems and curated database sources. For example, a metabolite such as prostaglandin E-2 identified using a Chemical Entities of Biological Interest identifier (ChEBI; (Hastings et al., 2016)) will be assigned additional, equivalent identifiers, and a standard name so that it has the same canonical form as an equivalent metabolite identified using a NCBI Medical Subject Heading identifier (MESH; **Figures S2A and S2B**). This procedure ensures that Agents in INDRA Statements take on canonical identifiers in multiple namespaces, irrespective of the identifier used in the original source of knowledge.

The final normalization procedure we performed was sequence normalization. This accounts for inconsistencies in attributed sequence positions of post-translational modifications, some of which involve outright errors in residue numbers, while others involve the implicit, interchangeable use of residue numbers between human and model organism reference sequences (Bachman et al., 2019). Commonly, human and mouse residue numbers are used interchangeably even through residue numbering in orthologous proteins frequently differs, so sequence normalization is necessary for accurate knowledge assembly.

After these steps were performed, Statements still containing ungrounded entities (∼38% of Statements contained Agents that lacked any identifiers) were filtered out, as were Statements containing non-canonical sequence positions (about 1% of Statements) as these likely arose from machine reading errors. Because the current study focuses on biology involving human genes, we also filtered the set of Statements to just those containing human genes and their families/complexes. Each of these processing and filtering steps operate at the level of individual Statements and change the overall number of Statements as well as proportion of Statements in the corpus from each input source, as shown in **Figure 2B**. The final corpus contained ∼2.9·10^6^ Statements after all filtering steps.

Once the normalization steps were complete, we used INDRA to combine Statements representing equivalent mechanisms from different sources into a single unique Statement; each unique Statement was associated with the supporting mentions from all contributing knowledge sources including curated databases and reading systems (**Figure 2C**). In some cases, multiple readers will have extracted the same mechanisms from the same sentence, but different reading systems often generated mentions supporting a specific Statement from different sentences in a given publications or even from different publications (**Figure 2C**). This highlights the substantial differences between reading systems and highlights the benefits of the multi-reader approach used in this paper. For the Benchmark Corpus, ∼2.9·10^6^ filtered Statements yielded ∼9·10^5^ unique Statements after combining duplicates (**Figure 2A**), with an average of ∼3 supporting mentions per Statement. However, the distribution of mentions per Statement was highly non-uniform, with a large number of Statements (63%) attributable to a single sentence or database entry, and a small number of Statements (82 in total) having >1,000 supporting mentions (**Figure 2D**). For example, the Statement that “*TP53 binds MDM2*” has 2,494 distinct pieces of evidence. Although noisy for high counts, the distribution of Statements having a given number of mentions appeared linear on a log-log plot (**Figure 2D**) implying a long-tailed distribution potentially following a power law. To confirm this, we fitted the observed mention distribution using two approaches: (i) linear regression of the complement cumulative distribution of mention counts on a log scale, which showed a strong linear relationship (r^2^=0.999, p<10^-17^), and implied a power law exponent of α,.2.33; and (ii) fitting directly to a power law using the *powerlaw* software package (Alstott et al., 2014), which showed that the distribution was fit by a power law with exponent α = 2.38 (standard error α= 0.0008) (**Figure 2E**) and was more likely than alternatives such as exponential (p < 10^-38^) or positive log-normal (p<10^-30^). Thus, the distribution of Statements having a given number of supporting mentions is similar to long-tailed distributions observed in a variety of domains including linguistics, computer networking and demographics (Clauset et al., 2009).

A significant benefit of jointly assembling mechanisms from both databases and literature is that curated interactions from databases become linked to textual evidence that support the interaction (**Figure 2F**). For example, the fact that RCHY1 ubiquitinates TP73 appears as a curated interaction in the NCI-PID database (Schaefer et al., 2009) with reference to PMID20615966 (Sayan et al., 2010), but without providing specific supporting text within that publication. In the Benchmark Corpus, INDRA aligns seven mentions obtained from text mining with the ubiquitination of TP73 by RCHY1 derived from four sentences in two more recent publications (Coppari et al., 2014; Wu et al., 2011) (**Figure 2F**). Such aggregation of evidence across curated databases and text mining systems is highly beneficial because it increases confidence in the accuracy and relevance of the mechanism. This is where INDRA, due to its automated nature, provides a substantial advantage for linking literature sources to specific interactions compared to comparable manual curation, which would be laborious and time consuming (Kemper et al., 2010).

### Detecting hierarchical relationships between mechanisms

Following processing, filtering, and the identification of duplicate Statements, the next assembly step is to identify relationships among “overlapping” Statements (**Figure 3A**). A pair of Statements is considered to be overlapping when one functions as a refinement (i.e., adds additional mechanistic detail) to the other. Although the analogy in this case is not perfect, something similar is required in genome assembly – if a shorter sequence is fully contained in a longer sequence, the shorter one is redundant. When such a relationship exists between two Statements, we say that the more detailed one “refines” the less detailed one. Refinement can happen at the level of entities (e.g., an Agent representing a protein family and another a specific member of that family), or molecular states and context (e.g., an explicit reference to a site of post-translational modification in one Statement and its omission in another). The refinement relationship between Statements is determined using a partial ordering logic that compares pairs of Statement based on their individual elements (where elements include the Agents involved in the Statement, and, depending on the type of Statement, post-translational modifications, cellular locations, types of molecular activity, etc.) and determines whether each element is either equivalent to or a refinement of the other (**Figure 3A**). To accomplish this, INDRA makes use of hierarchies of each relevant type of element, including proteins and their families and complexes drawn from FamPlex (Bachman et al., 2018), combined with chemical and bioprocess taxonomies from ChEBI and the Gene Ontology (Ashburner et al., 2000) (e.g., *MAP2K1* is a specific gene in the MEK family, **Figure 3A**, blue), protein activity types (e.g., kinase activity is a specific type of molecular activity, **Figure 3A**, red), post-translational modifications (e.g., phosphorylation is a type of modification, **Figure 3A**, green), and cellular locations (also obtained from the Gene Ontology; e.g., that the cytoplasm is a compartment of the cell, **Figure 3A**, purple). A Statement is also considered a refinement of another if it contains additional contextual details but is otherwise a match across corresponding elements. One example of such a refinement relationship is shown in **Figure 3B**, in which the first Statement (**Figure 3B**, top) describes an additional molecular state (*MAP2K1* being bound to *BRAF*) and mechanistic detail (T185 as the specific site of modification of *MAPK1*) over another Statement (**Figure 3B**, bottom) which omits these contextual details.

**Figure 3.**
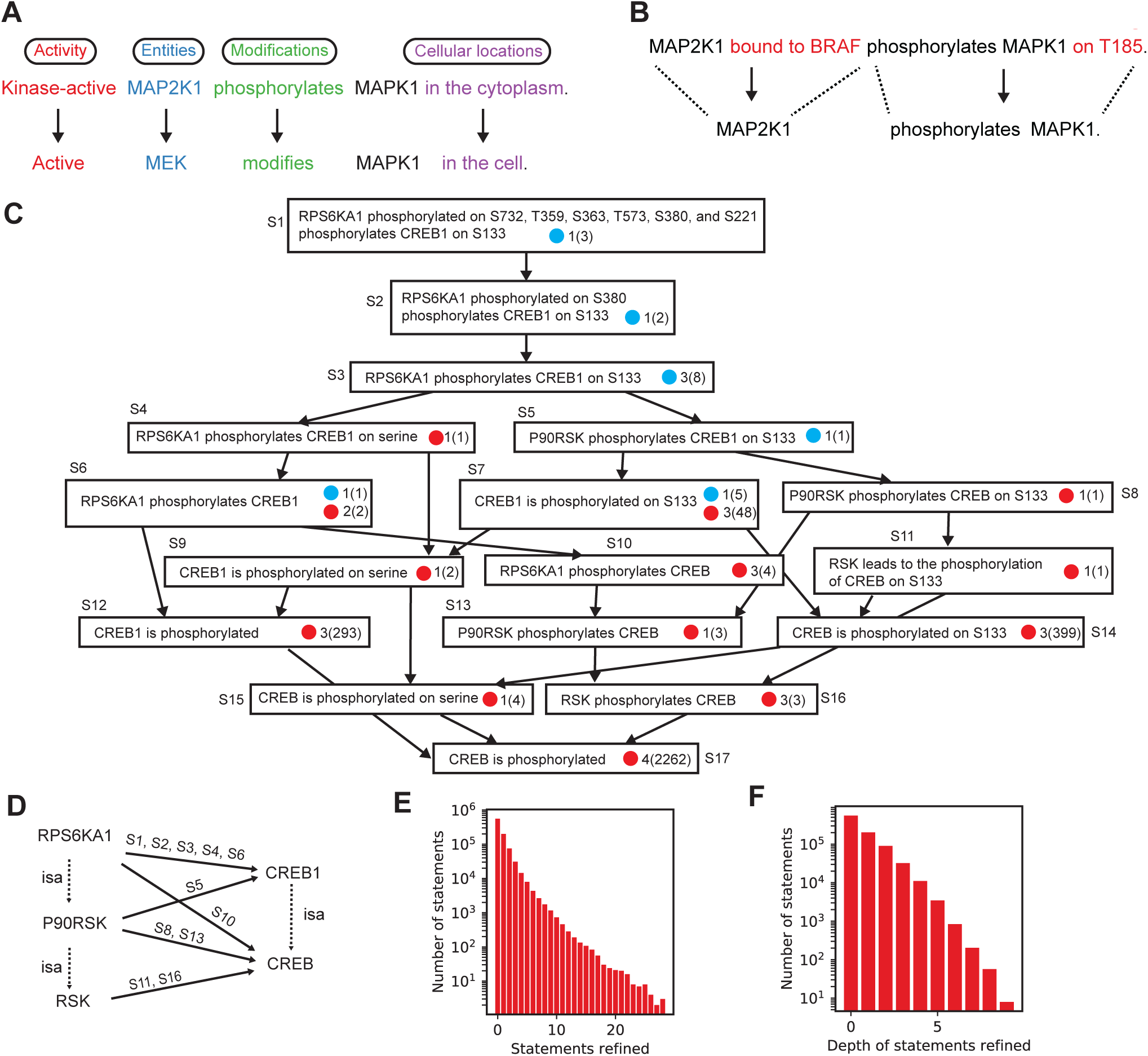
Identifying refinement relationships among Statements. (A) Refinement by hierarchies of Statement elements as defined by INDRA. The two Statements shown contain the same number and types of information but all elements in the top Statement are refinements of the corresponding elements in the bottom Statement according to the INDRA Statement hierarchies. (B) Refinement by additional context. The upper Statement contains all information in the lower one but also provides additional detail, making it a refinement of the one below. (C) Example refinement graph for a Statement from the example corpus. For clarity, the transitive reduction of the hierarchy is shown, and each Statement object is displayed via its English language equivalent. Each node in the graph represents a statement with blue or red circles representing evidence from pathway databases or mentions extracted by machine reading systems, respectively. Next to each blue or red circle, the number of different sources is shown with the overall number of mentions from these sources in parentheses. For example, the statement “*CREB1 is phosphorylated on S133*” has 5 pieces of evidence from one pathway database source, and 48 mentions extracted by three reading systems. Edges represent refinement relationships and point from more specific to less specific Statements. (D) Graph of family relationships (dotted *isa* edges) and Statements representing phosphorylation (solid edges, annotated with Statement identifiers from panel C), between different levels of specificity of the RSK and CREB protein families. (E) Number of Statements based on the total number of other Statements that they refine. (F) Number of Statements with different depths of Statements that they refine (i.e., the length of the longest path in the graph of refinement relations starting with the given Statement).

Pairwise refinement of relationships among Statements is most easily represented using a graph in which nodes represent Statements and directed edges point from a node representing a Statement to another node representing the Statement that it refines. Such Statement refinement graphs can be quite deep (i.e., the length of a directed path starting from a Statement can consist of a large number of edges going through many refined Statements). For example, the refinement subgraph for *RPS6KA1 phosphorylated on S732, T359, S363, T573, S380, and S221 phosphorylates CREB1 on S133* (**Figure 3C**, where *RPS6KA1* encodes the ribosomal S6 kinase and *CREB1* a transcription factor) has nine levels. The refinement relationships for this Statement reveal the varying levels of specificity at which a given mechanism is described in sources: *CREB is phosphorylated* has 2,268 mentions in the literature collected by 4 reading systems, *RPS6KA1 phosphorylates CREB1* has 3 mentions in total from both literature and curated databases, and *CREB1 is phosphorylated on S133* has 399 mentions. It is also worth noting that support from curated databases for these Statements (**Figure 3C**, blue circles) is not attributable to a single database source. For example the Statement labeled S1 in **Figure 3C** is derived only from Pathway Commons, S5 only from SIGNOR, and S7 only from HPRD (Mishra, 2006).

Organizing Statements hierarchically helps to ensure that an assembled model does not contain information that is mechanistically redundant. For instance, the Statements in **Figure 3C**, if viewed as a graph with nodes representing entities (RPS6KA1, CREB1, etc.) and edges representing phosphorylation reactions (**Figure 3D**, solid arrows) reveals five partially redundant edges (e.g., RPS6KA1→CREB1, P90RSK→CREB1, P90RKS→CREB, etc.) connecting members of the RSK and CREB protein families at different levels of specificity (e.g., P90RSK is a member of the RSK family, **Figure 3D**, dashed arrows). A key feature of INDRA is that it can recover Statement refinement relationships, enabling principled resolution of complex redundancies, for example, by retaining only Statements that are not refined by any other Statements (in the case of **Figure 3C**, the Statement labeled as S1 at the top of the graph). The refinement graph in **Figure 3C** also reveals how a highly specific Statement can serve as evidence for all the other Statements it subsumes, a relationship that is exploited when estimating Statement reliability.

We found that refinement relationships were common in the Benchmark Corpus: 38% of Statements refined at least one other Statement, and some Statements refined a large number of other Statements, including 89 Statements that refined at least 20 other Statements (**Figure 3E**). These Statements are typically ones that represent a canonical (i.e., often described) mechanism (for example, the mechanism by which members of the AKT protein family phosphorylate GSK3 proteins) at a high level of detail and subsume multiple variants of the same mechanism described at a lower level of detail. We also found that the Benchmark Corpus contained tens of thousands of refinements involving three or more levels (**Figure 3F**), emphasizing that many mechanisms across databases and literature are described at many levels of specificity. INDRA assembly can reconstruct these relationships and allow resolving the corresponding redundancy.

### Modeling the reliability of INDRA Statements with the help of a curated corpus

One of the most challenging problems in using mechanisms generated by text mining is the unknown reliability of the extracted information. While the notion of “reliability” includes conventional scientific concerns, such as the strength of a particular study or method (**Figure 4A**, upper left quadrant), in practice the overwhelming majority of incorrect assertions result from *technical* errors in machine reading (**Figure 4A**, lower left quadrant). Common reading errors include systematic misidentification of named entities, incorrect polarity assignment (e.g., classifying activation as inhibition), failure to recognize negative evidence (e.g., “A *does not* cause B’’), and difficulty distinguishing hypotheses from assertions and conclusions (e.g., “*we tested whether* A causes B” as opposed to “A causes B”) (Noriega-Atala et al., 2019; Valenzuela-Escárcega et al., 2018). These errors arise primarily because scientific text uses a wide range of non-standard naming conventions to refer to entities and uses complex grammatical structures to convey the confidence associated with a result or datapoint. Indeed, much of the art in scientific writing is to generate text that appears to progress inexorably from a hypothesis to the description of supporting evidence to a conclusion and its caveats. This type of writing can be difficult even for humans to fully understand. Addressing the technical errors of reading systems at the level of individual Statements is a prerequisite for addressing the additional issues that arise when Statements are combined into causal models (**Figure 4A**, right quadrants).

**Figure 4.**
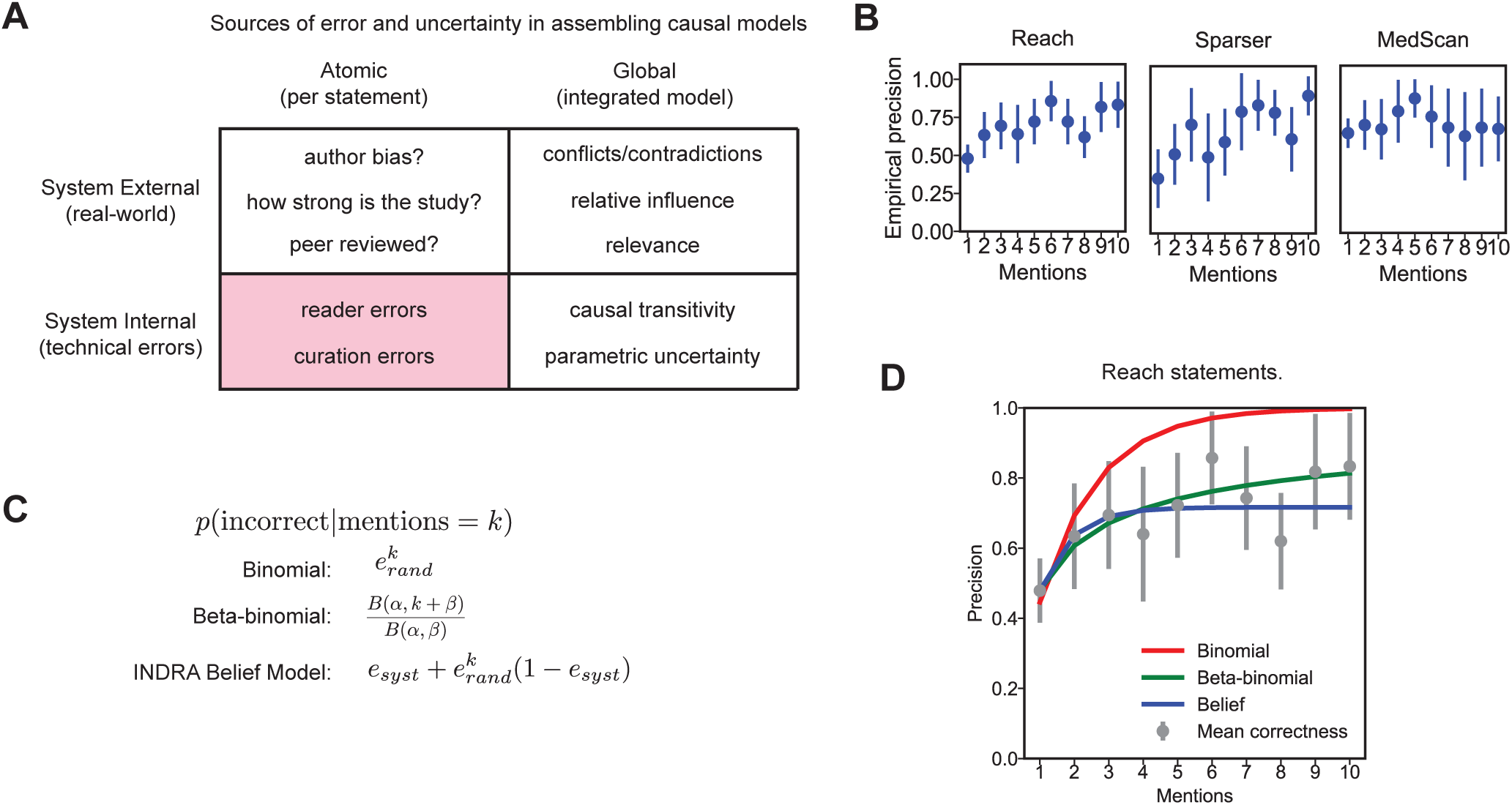
Estimating statement belief for a single machine reader. (A) A classification of sources of error and uncertainty in assembling causal models. Sources are classified according to whether they are external or internal to the INDRA system, and whether they arise at the level of individual Statements (atomic) or an integrated network or model (global). (B) Empirical precision of three reading systems based on the number of mentions supporting a given Statement extracted by that reader. (C) Mathematical formulas for Statement correctness for three different Belief Models. Each model specifies the probability that a Statement is incorrect overall given that a specific number *k* of mentions support it from a given source. *e_rand_*: random error for the source; *e_syst_*: systematic error for the source; *B(*Ill, β): Beta function. (D) Fits of the three belief models in (C) plotted against the empirical precision of Reach-extracted Statements.

To study the reliability of our assembled Statements, we sampled a set of Statements from the Benchmark Corpus. The sampled Statements had between 1 and 10 mentions per Statement and arose from five reading systems (Reach, Sparser, MedScan, RLIMS-P and TRIPS; we excluded the ISI/AMR system from this systematic analysis due to the low number of extractions it produced). Two of the authors, both of whom are PhD biomedical research scientists, used this to develop a Curated Corpus from the sampled Statements. Curation involved determining whether a given mention correctly supported a specific Statement based on human understanding of the sentence containing the mention and the overall context of the publication (see Methods). The resulting Curated Corpus data set covers ∼980 Statements with a combined total of ∼5,000 mentions (**Table 2**).

**Table 2:**
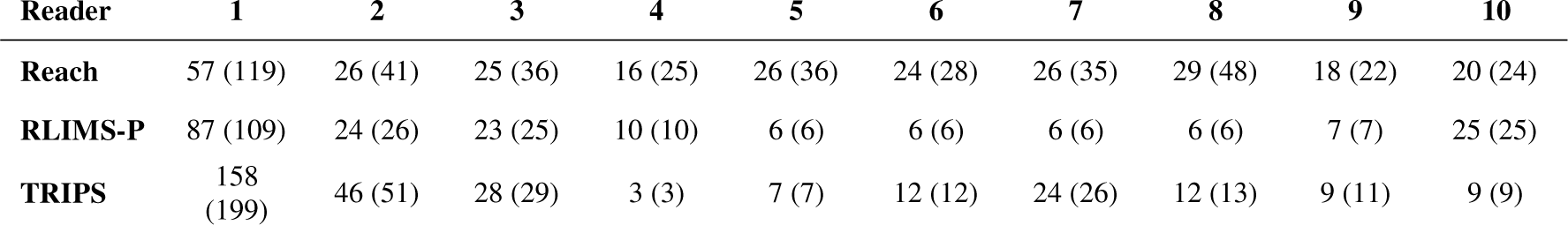
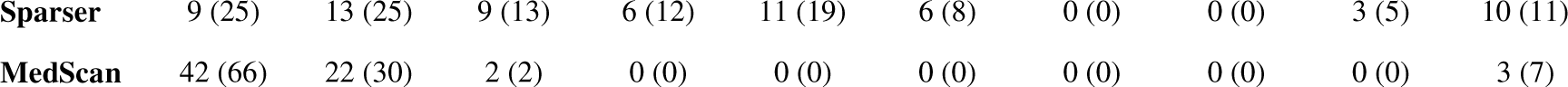
Summary of statement curation dataset. Entries are formatted as “number correct (total curated)”.

For a single reading system, the reliability of an extracted Statement has been observed to increase with the number of different supporting mentions (Valenzuela-Escárcega et al., 2018). We hypothesized that a Statement with multiple mentions would be more reliable if the supporting mentions had been independently extracted by more than one reading system. To test this idea, we used two complementary approaches to create models of Statement reliability: (i) structured probability models that build on empirical error characteristics of individual reading systems based on the Curated Corpus, and (ii) machine learning (ML) models trained on the Curated Corpus. Structured probability models require much less training data, however, machine learned models are generally more expressive and likely to be more accurate in predicting Statement reliability, given sufficient training data.

### Modeling the reliability of Statements from individual reading systems

We first examined the error characteristics of *individual* reading systems. For individual readers, analysis of the Curated Corpus showed that while Statements with more mentions are generally more reliable, in many cases Statements supported by many sentences were still incorrect due to the presence of systematic errors (**Figure 4B**). For example, the Sparser reading system extracted the Statement *MAOA binds MAOB* with ten mentions from ten different publications, but all extractions were incorrect because the system incorrectly interpreted “association” as referring to a physical interaction rather than a statistical association between MAOA and MAOB, which is what the original publications described. We compared three alternative probability models for their ability to capture the dependence of sentence reliability on mention count: (i) a simple binomial model, (ii) a beta-binomial model (a binomial model in which the probability of success at each trial follows a beta distribution), and (iii) a two-parameter model that captures both random and systematic errors – we termed this latter model the *INDRA Belief Model* (**Figure 4C**; see Methods). Each of the three models was independently fitted to data from the Curated Corpus using Markov chain Monte Carlo optimization (see Methods) (Foreman-Mackey et al., 2013). Both the beta-binomial model and the *INDRA Belief Model* outperformed the binomial model at predicting Statement correctness from mention counts, primarily due to their ability to capture the empirical observation that even high-mention Statements do not approach an accuracy of 100% (a phenomenon accounted for by modeling systematic reader errors) (**Figure 4D**, **Table 3**). The *INDRA Belief Model* performed slightly better than the beta-binomial model at predicting Statement correctness for both the Reach and Sparser reading systems (**Table 3)** due to its better fit to low mention-count Statements that make up the bulk of the corpus (**Figure 4D**, mentions 1, 2, and 3). An additional advantage of the *INDRA Belief Model* is that the random and systematic error rates *e_rand_* and *e_syst_*are interpretable and can be estimated heuristically by examining a small number of high-mention Statements (with precision approximately equal to *e_syst_*) and 1-mention Statements (with precision equal to *e_syst_* + (1 – *e_syst_*)*e_rand_*). This makes it possible to set reasonable parameters for the INDRA Belief Model based on prior intuition or examination of a small number of exemplary Statements. Since the *INDRA Belief Model* performed the best overall, it is used as the default model in INDRA when no curation data is available. However, we noted that the beta-binomial model more accurately fit the underlying distribution of correct mentions for each Statement, suggesting that further research is needed on such error models (**Figures S4A and S4B**).

**Table 3:** Maximum likelihood values for alternative belief models using best-fit parameters (lower values indicate a better fit).

| Model | Reach -log(Max<br>likelihood) | Reach -log(Max<br>likelihood) |
| --- | --- | --- |
| **Binomial (1 param)** | 366.2 | 120.31 |
| **Beta-binomial (2 params)** | 255.5 | 118.28 |
| **INDRA Belief Model (2 params)** | 254.6 | 118.24 |

### Multi-reader overlap is associated with higher Statement frequency and reliability

To better understand the potential for *multi*-reader reliability assessment, we characterized the extent of reader overlap in the Benchmark Corpus. We call two or more readers overlapping for a given Statement if they each produced mentions supporting that Statement. We found that 19% of assembled Statements had supporting mentions from two or more reading systems (**Table 4**; **Figures 5A and S5A**) but the bulk of Statements were supported exclusively by either Reach, Sparser, or MedScan (**Figure 5A**). The low overlap between readers is attributable to differences in their design, including their approaches to grammatical parsing, named entity recognition, associated resources (i.e., which lexical sources each reader incorporates), and the types of grammatical or semantic patterns that can be recognized. Low overlap among readers implies that using multiple reading systems in an integrated fashion via INDRA can increase coverage relatively to any single reading system.

**Figure 5.**
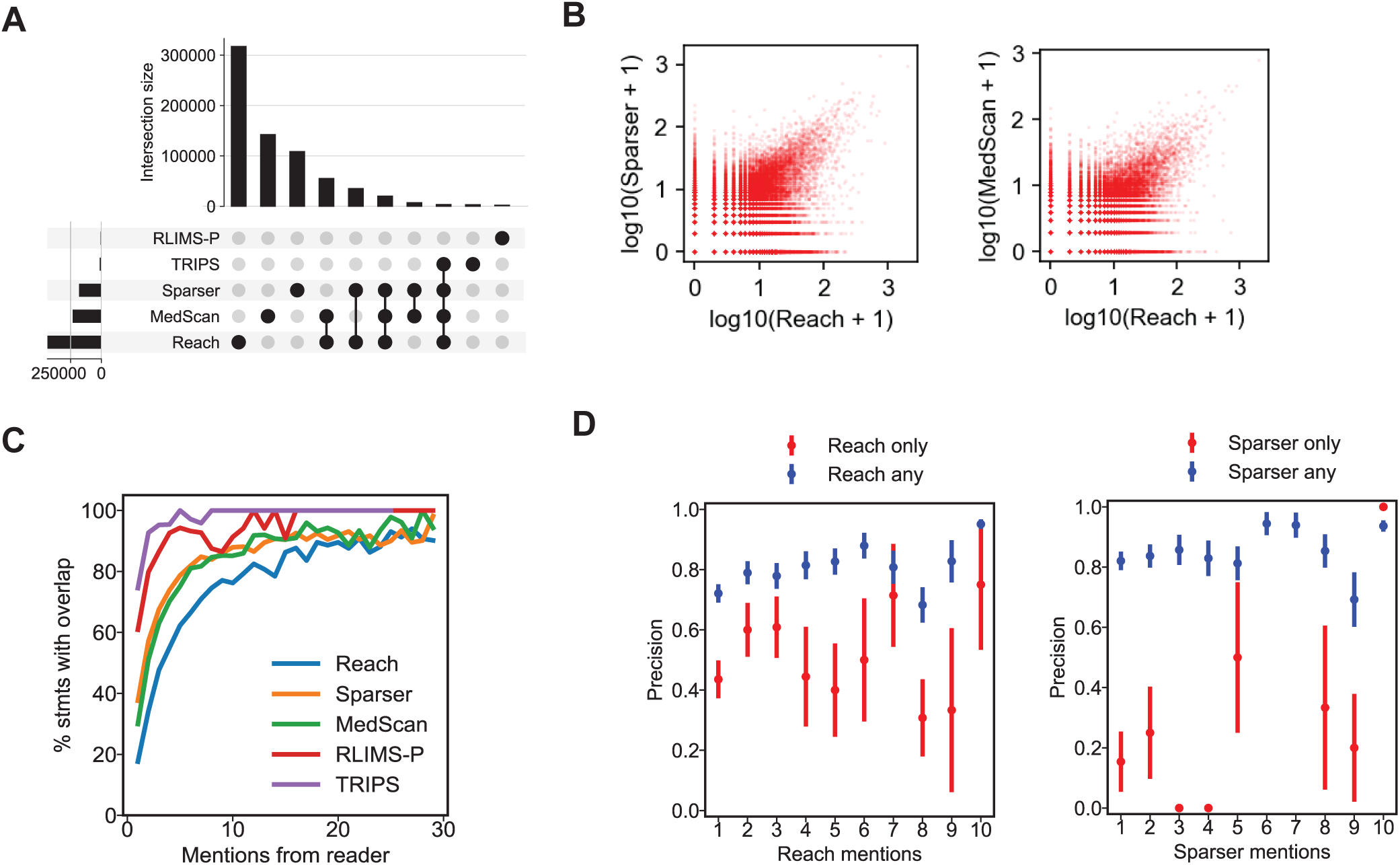
Estimating Statement belief with multiple machine readers combined. (A) Upset plot (equivalent to a Venn diagram with more than 3 sets) of Statement support for five machine reading systems integrated by INDRA. For a given Statement, two or more readers intersect if they each provide supporting mentions for it. (B) Number of mentions from Reach and Sparser (left) and Reach and MedScan (right) for a given Statement, each Statement being represented by a red dot. Mention counts are plotted on a logarithmic scale. (C) The percentage of Statements for which an intersection (i.e., any overlap) between reading systems is observed as a function of the number mentions from a given reader; the data are plotted separately for each of the five reading systems. (D) Empirical Statement precision as a function of the number of mentions from Reach (left) and Sparser (right), plotting the cases for which *only* Reach or Sparser provides supporting mentions for a Statement (red) and the case where all Statements are taken into account (blue).

**Table 4:**
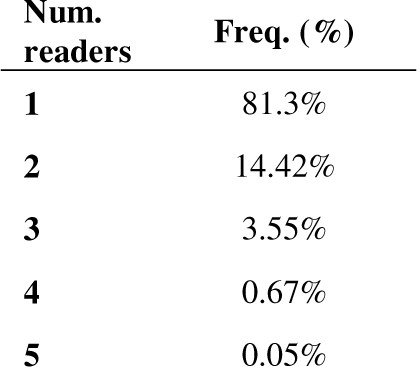
Frequencies of relations in corpus by total number of sources.

| Num.<br>readers | Freq. (%) |
| --- | --- |
| **1** | 81.3% |
| **2** | 14.42% |
| **3** | 3.55% |
| **4** | 0.67% |
| **5** | 0.05% |

Despite the relatively small overlap among readers, the number of mentions from each reader supporting a Statement showed substantial correlation, with both *ρ(Reach, Sparser)* and *ρ(Reach, MedScan)* > 0.6 (**Table 5**). We found, however, that these correlations in mention counts among Reach, Sparser, and MedScan were primarily driven by a subset of relations with very high numbers of mentions (**Figure 5B**). More generally, we found that reader overlap for a Statement increases as a function of the number of supporting mentions an *individual* reader extracted for the Statement (**Figure 5C**). Overall, these data support the observation that, if a mechanism represented by a Statement is described in many different sentences across input documents, multiple systems are likely to extract supporting mentions, and these will often come from different sentences and publications (as we showed in **Figures 2C** and **E**).

**Table 5:** Correlations between reader mention counts

|  | Sparser | MedScan | RLIMS-P | TRIPS |
| --- | --- | --- | --- | --- |
| **Reach** | 0.611 | 0.633 | 0.072 | 0.374 |
| **Sparser** |  | 0.454 | 0.114 | 0.420 |
| **MedScan** |  |  | 0.034 | 0.338 |
| **RLIMS-P** |  |  |  | 0.096 |

When we examined the relationship between reader overlap and Statement correctness using the Curated Corpus, we found that Statements supported by many mentions were more likely to overlap with other readers and be correct (**Figure S5B**, blue points along diagonals). Notably, in the case of Reach, the reader for which the most extensive subset of curated Statements was generated, we found that the probability of Statement correctness increased with the overall number of Reach mentions, but only for high-mention Statements that also included support from other readers (**Figure 5D**, blue points). For relations with support *only* from Reach, empirical correctness increased from 1 to 2 mentions (an observation consistent with the findings regarding the Reach system’s precision (Valenzuela-Escárcega et al., 2018)), but additional Reach-only mentions were not associated with substantial further increases in precision (**Figure 5D**, red points). Thus, in a multi-reader setting, the *absence* of reader overlap also plays a key role in assessing Statement reliability. These observations imply that combining multiple reading systems can be highly valuable when assessing Statement correctness based on supporting mentions. It also provides information that can be used by developers of reading systems to increase recall and precision.

### Two approaches to modeling the reliability of Statements from multiple readers

We evaluated two strategies for assessing the reliability of Statements using mention counts from multiple readers: (i) extending the INDRA Belief Model, and (ii) training machine learning models on the Curated Corpus. Even though reader errors were not in fact fully independent of each other (**Figure S5B**) we made an assumption of independence (Zhang, 2004) to extend the INDRA Belief Model to multiple reading systems while adding the fewest additional model parameters. Specifically, the model’s formulation of error estimates was changed to express the probability that all the mentions extracted by all the readers were *jointly* incorrect (see Methods). We compared the extended INDRA Belief Model to several different machine-learned classifiers for their ability to correctly predict Statement correctness based on mention counts from each reading system. Evaluated classifiers included Logistic Regression on log-transformed mention counts, k-Nearest Neighbors, support vector classifiers, and Random Forests (see Methods).

We found that, when mention counts were the only input feature, the INDRA Belief Model yielded the greatest area under the precision-recall curve (AUPRC), followed by the Logistic Regression and Random Forest models (**Table 6**, rows 1, 3, and 2, respectively). However, when the machine learning models were extended to make use of additional Statement features such as the Statement type, number of supporting articles (i.e., the number of distinct publications from which mentions were extracted), the average length of the mention texts (longer sentences were more likely to be incorrectly interpreted), and the presence of the word “promoter” in the sentence (a frequent indicator that a sentence describing a protein to DNA promoter interaction had been mis-extracted as a protein-protein interaction), they outperformed the INDRA Belief Model (Table 6, rows 8-13; see Methods). This implies that—as long as sufficient training data is available—machine-learned classifiers can use multiple features associated with Statements and their supporting mentions to boost performance as compared to the INDRA Belief Model which relies solely on mention counts.

**Table 6.** Comparison of belief models (using AUPRC as the metric) depending on model type, sources included, additional features taken into account, and whether more specific evidences are takin into account based on statement refinement relations. (Note: Statement is abbreviated as Stmt in the table).

| Group | Row | Model | Sources | Additional Features | More<br>specific<br>evidences | AUPRC |
| --- | --- | --- | --- | --- | --- | --- |
| Reader counts<br>only | 1 | **Belief Model** | **Readers** | N/A | **No** | **0.918** |
|  | 2 | Random Forest | Readers | None | No | 0.895 |
|  | 3 | Logistic Regression | Readers | None | No | 0.912 |
|  | 4 | k-NN | Readers | None | No | 0.896 |
|  | 5 | SVC | Readers | None | No | 0.87 |
| More features | 6 | Random Forest | Readers | Stmt type, # PMIDs | No | 0.912 |
|  | 7 | Random Forest | Readers | Stmt type, # PMIDs, Avg.<br>evidence len. | No | 0.918 |
|  | 8 | Random Forest | Readers | Stmt type, # PMIDs, "promoter" | No | 0.923 |
|  | 9 | **Random Forest** | **Readers** | **Stmt type, # PMIDs, "promoter", Avg. evidence len.** | **No** | **0.927** |
|  | 10 | Logistic Regression | Readers | Stmt type, # PMIDs | No | 0.923 |
|  | 11 | Logistic Regression | Readers | Stmt type, # PMIDs, Avg. evidence len. | No | 0.923 |
|  | 12 | Logistic Regression | Readers | Stmt type, # PMIDs, "promoter" | No | 0.924 |
|  | 13 | Logistic Regression | Readers | Stmt type, # PMIDs, "promoter", Avg. evidence len. | No | 0.924 |
|  | 14 | k-NN | Readers | Stmt type, # PMIDs | No | 0.898 |
|  | 15 | SVC | Readers | Stmt type, # PMIDs | No | 0.879 |
| Reader counts only, specific evidences | 16 | **Belief Model** | **Readers** | N/A | **Yes** | **0.919** |
|  | 17 | Random Forest | Readers | None | Yes | 0.913 |
|  | 18 | Logistic Regression | Readers | None | Yes | 0.914 |
|  | 19 | k-NN | Readers | None | Yes | 0.897 |
|  | 20 | SVC | Readers | None | Yes | 0.888 |
| More features, specific evidences | 21 | Random Forest | Readers | Stmt type, # PMIDs | Yes | 0.925 |
|  | 22 | Random Forest | Readers | Stmt type, # PMIDs, Avg. evidence len. | Yes | 0.928 |
|  | 23 | Random Forest | Readers | Stmt type, # PMIDs, "promoter" | Yes | 0.931 |
|  | 24 | **Random Forest** | **Readers** | **Stmt type, # PMIDs, "promoter", Avg. evidence len.** | **Yes** | **0.933** |
|  | 25 | Logistic Regression | Readers | Stmt type, # PMIDs | Yes | 0.924 |
|  | 26 | Logistic Regression | Readers | Stmt type, # PMIDs, Avg. evidence len. | Yes | 0.924 |
|  | 27 | Logistic Regression | Readers | Stmt type, # PMIDs, "promoter" | Yes | 0.925 |
|  | 28 | Logistic Regression | Readers | Stmt type, # PMIDs, "promoter", Avg. evidence len. | Yes | 0.925 |
|  | 29 | k-NN | Readers | Stmt type, # PMIDs | Yes | 0.892 |
|  | 30 | SVC | Readers | Stmt type, # PMIDs | Yes | 0.9 |
| All sources, more features, specific evidences | 31 | Belief Model | Readers, DBs | N/A | Yes | 0.927 |
|  | 32 | Random Forest | Readers, DBs | None | Yes | 0.933 |
|  | 33 | Random Forest | Readers, DBs | Stmt type, # PMIDs | Yes | 0.935 |
|  | 34 | Random Forest | Readers, DBs | Stmt type, # PMIDs, Avg. evidence len. | Yes | 0.936 |
|  | 35 | Random Forest | Readers, DBs | Stmt type, # PMIDs, "promoter" | Yes | 0.94 |
|  | 36 | **Random Forest** | **Readers, DBs** | **Stmt type, # PMIDs, "promoter", Avg. evidence len.** | **Yes** | **0.941** |
|  | 37 | Logistic Regression | Readers, DBs | None | Yes | 0.925 |
|  | 38 | Logistic Regression | Readers, DBs | Stmt type, # PMIDs | Yes | 0.934 |
|  | 39 | Logistic Regression | Readers, DBs | Stmt type, # PMIDs, Avg. evidence len. | Yes | 0.934 |
|  | 40 | Logistic Regression | Readers, DBs | Stmt type, # PMIDs, "promoter" | Yes | 0.935 |
|  | 41 | Logistic Regression | Readers, DBs | Stmt type, # PMIDs, "promoter", Avg. evidence len. | Yes | 0.936 |

The ability of INDRA to identify refinement relationships among Statements (**Figure 3**) has the added benefit that it allows mentions to be combined across different levels of detail for use in reliability estimation. For example, any evidence for the specific Statement “*MAP2K1 phosphorylates MAPK1 on T185*” also supports the more generic Statement “*MEK phosphorylates ERK*.” We found that combining more specific mentions with mentions directly associated with a specific Statement improved precision and recall: the AUPRC of the Random Forest model increased from 0.895 to 0.913 when using only mention counts (**Table 6**, row 2 vs. 17), and from 0.927 to 0.933 when using all features (**Table 6**, row 9 vs. 24). Further, when we examined the effect of incorporating overlapping mentions from curated databases as features alongside mentions from readers, we found that the Random Forest model’s AUPRC increased to 0.941 – the highest AUPRC reached across all models and conditions.

Because readers perform differently on the same input text, Statements supported by multiple readers are less common than Statements supported by a single reader but our analysis showed that both the existence of reader overlap as well as lack of overlap for a given Statement can be informative for predicting Statement correctness. Moreover, in the absence of human-curated data across multiple Statement features – a type of data that is laborious to generate – a parametric model (such as the INDRA Belief Model) based on the error profiles of individual readers can perform well from a precision-recall perspective. When sufficient curated training data is available, machine learning models such as Random Forests can achieve greater performance, obtaining an AUPRC greater than 0.9 in several different configurations. These findings provide empirical support for INDRA’s approach to assembling sets of Statements from multiple text mining and curated database sources with principled estimates of correctness. Both the INDRA Belief Model and the machine-learned classifier models are available in the *belief* submodule of INDRA and allow parameters to be either manually set or estimated from curation data.

### Validation of assembled mechanisms and comparison against curated resources

To test INDRA on a prototypical biocuration task, we compared the subset of Statements representing human protein-protein interactions (PPIs) in the Benchmark Corpus to the BioGRID database (Oughtred et al., 2019). BioGRID is a curated public database containing structured information on protein-protein and protein-small molecule interactions, as well as genetic interactions obtained from multiple organisms. These interactions were extracted by expert curators from a combination of high-throughput datasets and focused studies. As a measure of the utility of INDRA for biocuration we determined (i) the number of previously-uncurated PPIs that the INDRA Benchmark Corpus could add to BioGRID and (ii) the amount of new literature evidence that it could added to PPIs currently in BioGRID. We used our best-performing Random Forest model to assign a belief to each INDRA Statement in the Benchmark Corpus.

The Benchmark Corpus contained ∼26,000 Statements representing PPIs already in BioGRID, and ∼101,000 PPIs that were absent (**Figure 6A**); the latter potentially represent known but previously uncurated interactions. Grouping all PPIs in bins defined by belief score, we found that belief score was highly correlated with the likelihood of a PPI being curated in BioGRID (**Figure 6B**). This provides a quantitative corroboration of the belief scores and, by extension, suggests that a substantial number of the potentially new PPIs involve reading errors which are accounted for by low belief scores. Because the belief scores obtained from the Random Forest model can be interpreted as calibrated probabilities of correctness, they can be used to estimate the *number* of Statements in each bin that are expected to be correct. The proportion of Statements in BioGRID was consistently below the belief score for the bin, suggesting that each bin contained correctly extracted but uncurated PPIs (**Figure 6B**, blue line below diagonal). Conservatively assuming that all Statements found in BioGRID were correctly extracted, we estimated a lower bound of 28,600 correct but uncurated PPIs in the Benchmark Corpus, a 6% increase over the ∼480,000 unique human PPIs in BioGRID.

**Figure 6.**
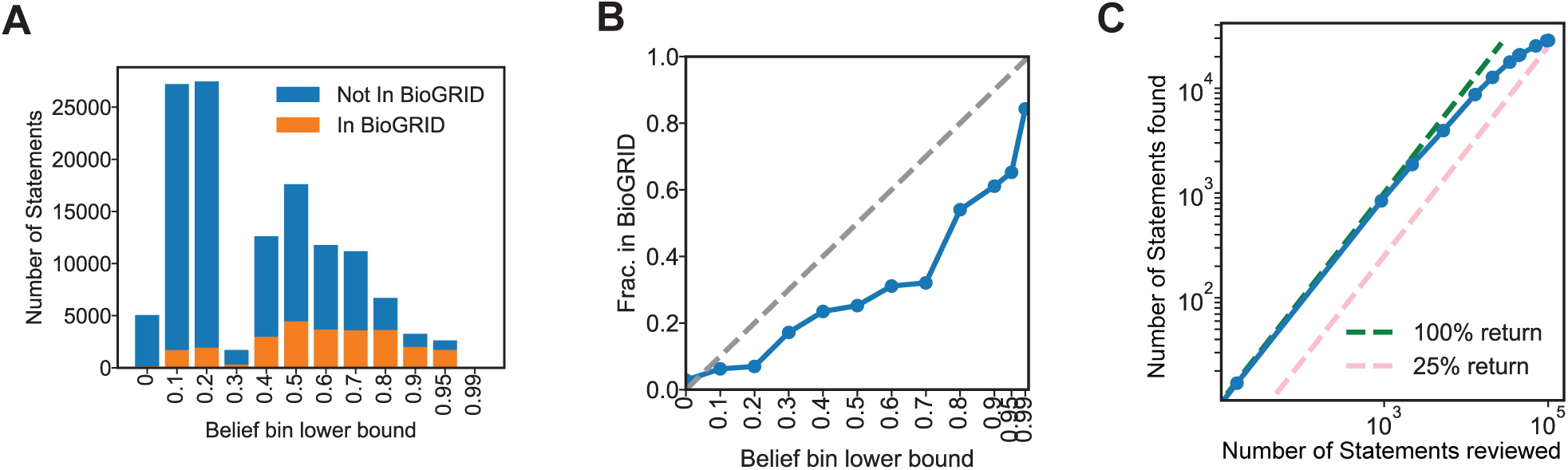
Comparison of INDRA-assembled mechanisms with a curated resource, BioGRID. (A) Number of INDRA Statements representing PPIs (i.e., complex formation between two human proteins) grouped into bins by their belief score (as determined by a random forest belief model), differentiating whether the PPI represented by the Statement appears in BioGRID (orange) or not (blue). (B) Fraction of INDRA Statements representing PPIs that appear in BioGRID grouped into bins by their belief score. A gray dashed line shows the expected fraction of correct Statements in each belief bin. The space between the gray and blue lines (i.e., between the expected fraction of correct Statements in each bin and the fraction of Statements that appear in BioGRID) represents an estimate of the set of correct Statements missing from BioGRID. (C) Plot showing estimated curation yield if Statements were reviewed by decreasing belief score for inclusion into a curated resource. The blue line plots the number of correct Statements expected to be found as a function of the number of Statements reviewed, with green and pink dashed lines serving as guides showing 100% return (i.e., every reviewed Statement is correct) and 25% return (i.e., 1 out of 4 reviewed Statements is correct).

As a practical matter, correct and incorrect Statements could be efficiently separated by manual curation, focusing first on Statements with the highest belief scores. The ∼2,200 uncurated Statements with belief scores > 0.9 would be expected to yield >1,870 PPIs, or roughly six correct for every seven reviewed. Statements with lower belief scores are more numerous but also have a lower expected yield: 18,700 correct but uncurated Statements would be expected among the the 41,600 Statements with belief scores between 0.4 and 0.9, with the curation yield starting at 67% (for Statements with belief between 0.8 and 0.9) to 29% (for Statements with belief between 0.4 and 0.5) (**Figure 6C**). By way of illustration, we examined one PPI not currently in BioGRID that involved binding of the KIF1C kinesin to RAB6A, a GTPase and regulator of membrane trafficking. INDRA assembled a total of 40 mentions supporting this PPI, extracted by two machine reading systems (Reach and Sparser), into a Statement with belief score 0.82. Human curation confirmed that the interaction had been reliably demonstrated using both co-immunoprecipitation and reconstitution experiments (Lee et al., 2015).

A second application of INDRA is to add evidence for PPIs already in BioGRID and thereby (i) provide new and different types of evidence for an existing PPI (e.g., mass spectrometry vs. 2-hybrid interaction), (ii) reveal additional biological settings or cell types in which a PPI might occur, and (iii) provide additional mechanistic detail about a particular PPI. As an example of (i) and (ii), BioGRID lists only three publications as a reference for the interaction between brain-derived neurotrophic factor (BDNF) and the NTRK2 receptor tyrosine kinase, whereas the INDRA Benchmark Corpus contains 168 mentions of this interaction from a total of 94 publications. Some of these additional publications provide primary experimental evidence for this interaction (e.g., (Vermehren-Schmaedick et al., 2014) and (Wang et al., 2009a)) discuss the role of the BDNF-NTRK2 interaction in important biological or clinical settings. As an example of (iii), the interaction between paxillin (PXN) and the tyrosine kinase PTK2B is supported by six references in BioGRID; INDRA not only identified 49 mentions from 18 different publications supporting this PPI, but assembled a Statement with substantially more mechanistic information: namely that PTK2B, when phosphorylated on Y402, phosphorylates PXN on Y118 (Moody et al., 2012; Park et al., 2006). This example shows that for a PPI lacking mechanistic detail, INDRA can illuminate the directionality and type of regulation, as well as the amino acids involved in posttranslational modifications.

### Detecting and explaining gene dependency correlations with an assembled causal network

To study how networks that incorporate text-mined information can aid in the interpretation of functional genomic datasets, we used INDRA to detect and explain significant gene dependencies in the Cancer Dependency Map (https://depmap.org) (Meyers et al., 2017; Tsherniak et al., 2017). The DepMap reports the effects of RNAi or CRISPR-Cas9 mediated gene inactivation on cell viability and growth in >700 cancer cell lines using a competition assay. In this assay, the effect of gene inactivation is assessed by determining the rate at which a specific knockout (or knockdown) disappears from a co-culture comprising cells transfected with a genome-scale RNAi or CRISPR-Cas9 library. It has previously been observed that that genes whose knockouts have similar effects on viability across a large number of cell lines —a phenomenon known as codependency—frequently participate in the same protein complex or pathway (Doherty et al., 2021; Meyers et al., 2017; Pan et al., 2018; Rahman et al., 2021; Shimada et al., 2021; Tsherniak et al., 2017). For example, CHEK2 and CDKN1A have a correlation coefficient of 0.359 and 0.375 in DepMap CRISPR and RNAi data, respectively (**Figure 7A**), and this codependency can be explained by the fact that the CHEK2 kinase is an activator of CDKN1A (also known as p21) and that the two genes jointly regulate cell cycle progression. To obtain robust measures of gene co-dependencies, we combined the CRISPR and RNAi perturbation data by converting the Pearson correlation coefficients for each gene pair into signed z-scores and computing the combined z-score between the two datasets using Stouffer’s method (**Figure 7A**). In analyzing the data, we first accounted for a bias also observed by others (Dempster et al., 2019; Rahman et al., 2021), namely that many of the strongest correlations are between mitochondrial genes (**Figure 7B).** These correlations have been described as an artifact of the screening method (such as the timepoint of the viability measurements relative to cell doubling time) rather than reflecting true co-dependencies (Rahman et al., 2021). We considered the correlations among these genes to be “explained” *a priori* due to their shared mitochondrial function, and using the mitochondrial gene database MitoCarta as a reference (Rath et al., 2021), we excluded correlations among them from subsequent analysis.

**Figure 7.**
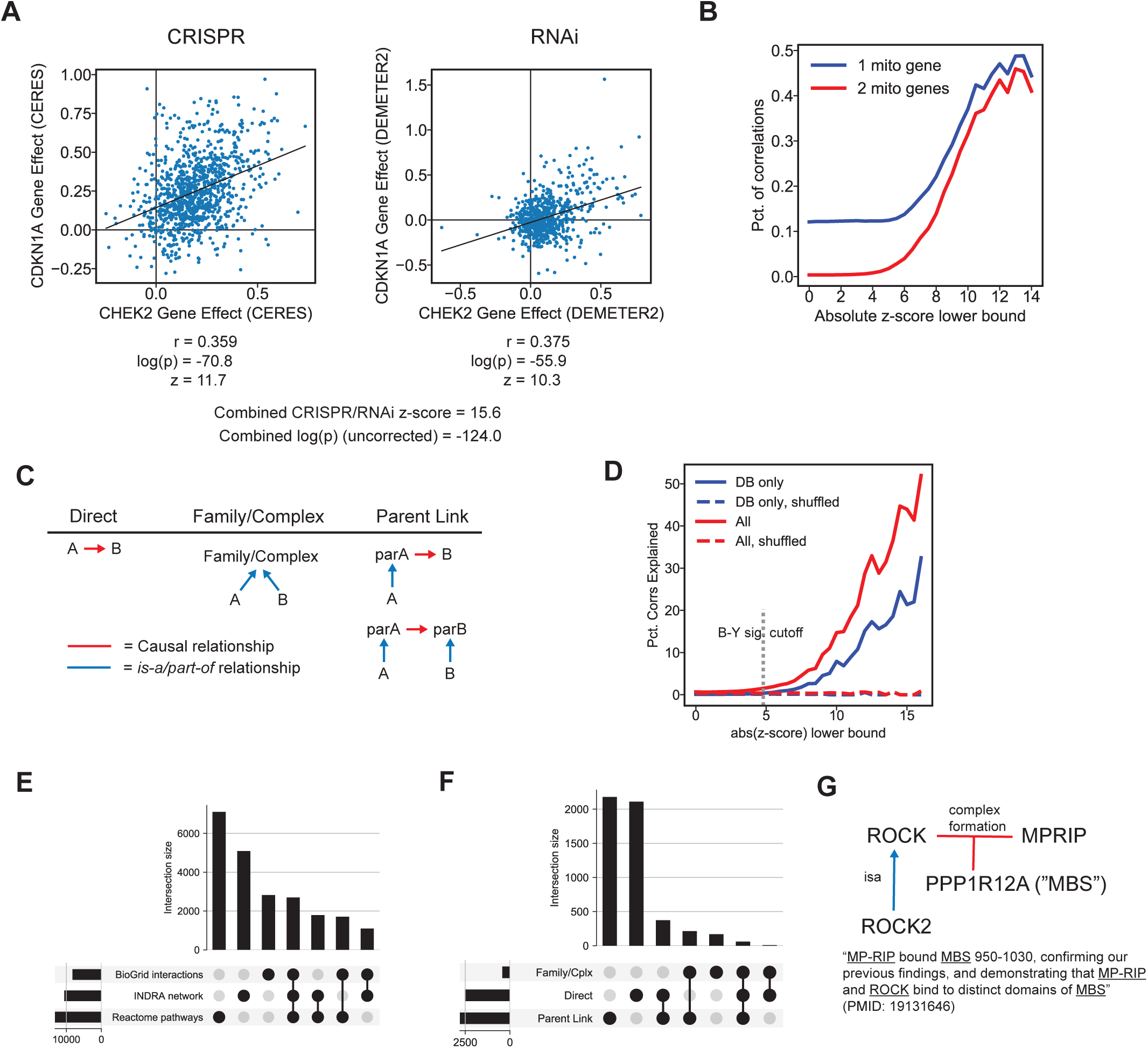
Detecting and explaining gene codependency in cancer cell lines using an INDRA-assembled network. (A) CRISPR (left) and RNAi (right) data from DepMap showing the codependency of the CHEK2 and CDKN1A genes across a panel of cancer cell lines (each blue dot represents a cell line, placed according to normalized cell viability change upon gene perturbation). Black lines show the linear regression plot over the cell line viability values. (B) Percent of gene codependencies (i.e., correlations) involving one or two mitochondrial genes as a function of the absolute z-score corresponding to the codependency. (C) Patterns of network nodes and edges that constitute an “explanation” for an observed DepMap codependency, including “Direct” (a direct edge between two specific genes A and B), “Family/Complex” (two genes A and B are part of the same family or complex), and “Parent Link” (where one or both of the specific genes A and B are related via a parent family/complex they are part of). (D) Percent of codependencies/correlations explained using the INDRA network when considering all edges (red) or only edges supported by curated databases, excluding text mining (blue), with randomly shuffled controls shown. (E) Upset plot showing the intersection of explanations for DepMap codependencies provided by three networks: BioGRID interactions, the INDRA network, and Reactome pathways. (F) Upset plot showing the intersection of three types of explanation for DepMap codependencies provided by the INDRA Network, corresponding to explanation patterns shown in panel D. (G) An example explanation for the codependency between ROCK2 and MPRIP derived from the INDRA network. INDRA provides evidence for a complex in which ROCK (the protein family of which ROCK2 is a member) binds MPRIP in a three-way complex with PPP1R12A (also called MBS) through the mention shown at the bottom (extracted from Wang et al, 2009).

From the Benchmark Corpus of assembled INDRA Statements, we generated a network model in which each node represents a human gene and each directed edge corresponds to an INDRA Statement (such as Phosphorylation, Activation, etc.) connecting two nodes. We used the resulting network for two tasks: first, to constrain the number of hypotheses tested when determining the statistical significance of codependency correlations, and second, to find mechanistic explanations for the observed codependencies. For the first task, we calculated the number of codependencies that were significant at a false discovery rate (FDR) of 0.05 using three methods for controlling FDR with and without the use of the network to limit the number of hypotheses tested (**Table 7**). Overall, fewer codependencies were identified as significant when we restricted comparisons to relationships in the INDRA-assembled network, both because the network is incomplete and because many codependencies reflect indirect functional relationships not captured by a single direct edge in the network. However, many codependencies (4,007 using Benjamini-Yekutieli FDR correction) were detected as significant *only* when using the network (**Table 7**, “INDRA only”) due to the smaller number of hypotheses tested. Moreover, the majority of these (2,729) were based on interactions obtained only from machine reading, of which >60% were supported by a Statement with a belief score greater than 0.5.

**Table 7.** Number of codependencies detected at a significance cutoff of p < 0.05 without multiple hypothesis correction or after one of three methods for multiple hypothesis testing correction (Bonferroni, Benjamini-Hochberg, and Benjamini- Yekutieli). Results are shown for a case in which no prior is used and data is analyzed directly (“No prior”), or when an INDRA prior is used (“INDRA prior / Total”). The rightmost column shows the number of novel codependencies recovered exclusively when an INDRA prior was used along with correction for multiple testing (“INDRA prior / INDRA only”). †Figures for uncorrected p-values do not apply to the “INDRA prior / INDRA-only” case because without correction for multiple testing, the prior does not play a role in determining significance. Figures are shown for the “INDRA prior / Total” case to establish the number of codependencies with uncorrected p-values > 0.05 that fall within the scope of the INDRA network; this serves as an upper bound for the number of correlations determined to be significant with the different approaches to multiple testing shown in the bottom three rows.

|  | No prior | INDRA prior |  |
| --- | --- | --- | --- |
|  |  | Total | INDRA only |
| Number of comparisons (non-mitochondrial) | 121,778,711 | 265,874 | N/A† |
| Correlations with uncorrected $p < 0.05$ | 21,526,511 | 63,926† | N/A† |
| Significant corrs after Bonferroni | 99,544 | 4,982 | 1,836 |
| Significant corrs after Benjamini-Hochberg | 5,025,535 | 30,127 | 7,506 |
| Significant corrs after Benjamini-Yekutieli | 972,831 | 12,812 | 4,007 |

Conversely, the existence of a codependency added context to text-mined mechanisms. For example, the negative correlation between ERBB2 and STMN1 (ρ= −0.146 in DepMap CRISPR data) was associated with a single INDRA phosphorylation Statement in the Benchmark Corpus; the fact that the codependency correlation is *negative* indicates that ERBB2 phosphorylation of STMN1 inhibits it (a finding corroborated by (Benseddik et al., 2013)). Similarly, the negative correlation between GRB10 and IRS2 (ρ=-0.137 in CRISPR) is consistent with reports that the binding of GRB10 to IRS2 is inhibitory. This provides context for the INDRA Statement derived from (Keegan et al., 2018; Mori et al., 2005) that “*GRB10 binds IRS2*” and is particularly interesting because the effect of GRB10 binding to IRS2 has been reported as both inhibitory (Wick et al., 2003) and activating (Deng et al., 2003). The negative DepMap correlation suggests that the inhibitory effect is more relevant in the context of the two genes’ co-regulation of cell viability. Overall, these findings suggest that an INDRA-assembled networks can lead to the detection of codependencies that would otherwise be missed, and—as the previous two examples show—the combined information from data and assembled knowledge provides deeper mechanistic insight into each interaction than data alone.

In addition, we tested whether the Benchmark Corpus could provide mechanistic explanations of DepMap codependencies beyond what can be explained by curated pathway databases. We considered three types of relationships to be explanatory: (i) direct causal relationships where one gene was reported to regulate, modify or interact with another, e.g. the inhibition of TP53 by MDM2, (**Figure 7C**, “Direct”), (ii) information that the two correlated genes were members of the same protein family or complex, as indicated by FamPlex relations (Bachman et al., 2018) (**Figure 7C**, “Family/Complex”), or (ii) a link between the parent family/complex of a gene and another gene or its parent family/complex (**Figure 7C**, “Parent Link”). To measure the impact of text mining, we derived a smaller, “database-only” network from the Benchmark Corpus by excluding edges that were supported *only* by text mining evidence from the “full” network. As a control, we permuted the node labels of both the full and database-only networks and repeated our analysis. We found that the full network explained a greater proportion of codependencies than the database-only network (22% vs. 11% for codependencies with |z-score| > 6), with similar improvements at all significance levels (**Figure 7D**). This improvement is striking considering the text mining results were drawn from a corpus that constitutes only a fraction of what is currently available in PubMed. We also found that for either network, stronger codependencies were more likely to be explainable than weaker ones (**Figure 7D**), highlighting that the curated and published mechanistic knowledge (that is likely to be picked up by INDRA) is generally biased towards the most robust functional relationships.

To better characterize how INDRA-assembled networks provide mechanistic context for relationships in DepMap, we compared codependencies explainable via the full INDRA network to those explainable via interactions in BioGRID or by co-membership in a Reactome pathway. Of the 345,077 non-mitochondrial gene pairs with DepMap codependency correlations above the Benjamini-Yekutieli significance cutoff, only 21,475, or 6.2%, could be explained by BioGRID interactions, a common Reactome pathway, or the INDRA network, highlighting the many potential functional relationships in DepMap without a literature precedent. Membership in a common Reactome pathway, the least specific type of explanation, accounted for the largest number of explanations, including 6,952 codependencies explainable only via this information (**Figure 7E**). The INDRA network accounted for the next-highest number of unique explanations with 4,819 (**Figure 7E**). Interestingly, a majority of these were attributable to regulatory relationships mediated by families and complexes of which specific codependent genes were members (**Figure 7F**, “Parent Link” explanations). While less stringent than explicit gene-gene relationships, these family-mediated connections can produce compelling explanations for genes commonly described at the level of families and complexes (Bachman et al., 2018). For example, the strong negative correlation between MPRIP and ROCK2 (ρ=-0.291) is explained by multiple text mined Statements linking MPRIP to the ROCK protein family (referred to generically as “ROCK” or “Rho kinase”) via their joint binding to the myosin-binding subunit of the myosin light chain phosphatase (gene PPP1R12A, **Figure 7G**) (Nunes et al., 2010; Surks et al., 2003; Wang et al., 2009b).

The remainder of the INDRA-dependent explanations were derived from Statements involving two specific codependent genes (**Figure 7F**, “A->B”). While these explanations are “direct” in the sense that two genes are linked by an edge in the INDRA network, the relationships may not involve physical binding and might therefore have intermediaries (a mechanistically indirect connection). Such indirect mechanisms can be an advantage in many systematic explanation tasks. For example, the strong correlation between BRAF and MITF (ρ=0.456) cannot be explained by a common Reactome pathway, a physical interaction in BioGRID, or interactions in any of the pathway databases incorporated in the INDRA network. However, BRAF and MITF are linked by an INDRA network edge derived from 20 text-mined Statements (supported by 59 distinct mentions) characterizing their complex mutual regulatory relationship. The Statements correctly capture the evidence that *oncogenic* BRAF *activates* the expression of MITF through the transcription factor BRN2 (Kumar et al., 2014) whereas *wild type* BRAF in melanocytes *inhibits* MITF expression due to the lack of expression of BRN2 (Wellbrock et al., 2008). Because INDRA can represent molecular states on Agents (in this case BRAF vs. its mutated form BRAF-V600E) these extracted Statements are able to provide machine-readable information differentiating the two distinct contexts. Finally, we noted that interactions obtained exclusively from text mining were not restricted to well characterized or indirect relationships: for example, the INDRA network also incorporates a Statement extracted from a single sentence explaining the correlation between DOCK5 and BCAR1 (better known as p130Cas) as arising from their joint interaction with the scaffold protein CRK (Frank et al., 2017). Despite their robust correlation (ρ=0.361), DOCK5 and BCAR1/p130Cas have only been co-mentioned in a total of three publications in PubMed.

## DISCUSSION

In this paper, we described a method, implemented in INDRA software, for robust, automated assembly of mechanistic causal knowledge about biological interactions. The method normalizes information from heterogeneous sources, including both curated databases and text mining systems, and integrates this information by identifying relationships between Statements and using statistical models to estimate the reliability of each Statement based on the totality of the supporting evidence. The corpus used in this paper (∼570,000 articles) covers only a fraction of the published biomedical literature (>30 million articles). Nevertheless, we demonstrate that it is possible to meaningfully extend curated interaction databases and provide explanations for gene dependency correlations in the Cancer Dependency Map. INDRA enriches biocuration and data analysis efforts in three ways, i) by aggregating and normalizing new, previously uncurated mechanisms directly from the literature in machine-readable form, ii) by adding mechanistic detail (activation, modification, binding, etc.) to generic PPIs or empirical relationships, and iii) by supplying supporting evidence and context from the literature. Others can make use of INDRA tools since they are open source (https://github.com/sorgerlab/indra) and well-documented (https://indra.readthedocs.io). INDRA has already been used for diverse knowledge assembly, curation, and analysis tasks, including network-based gene function enrichment (Ietswaart et al., 2021), causal analysis of viral pathogenesis (Zucker et al., 2021), drug target prioritization for acute myeloid leukemia (Wooten et al., 2021), assembling knowledge about protein kinases (Moret et al., 2021), assisting manual biocuration efforts (Glavaški and Velicki, 2021; Hoyt et al., 2019a; Ostaszewski et al., 2021), and helping authors capture mechanistic findings in computable form (Wong et al., 2021).

The method described here is related to prior work on the integration of biological databases (Rodchenkov et al., 2020; Szklarczyk et al., 2021; Türei et al., 2016), assembly of biological knowledge graphs (Himmelstein et al., 2017; Hoyt et al., 2019b), large-scale biomedical event extraction (Van Landeghem et al., 2011), and estimation of the reliability of individual interactions in knowledge graphs (Jia et al., 2019; Neil et al., 2018). However, it goes beyond the straightforward aggregation of interactions from multiple sources by 1) systematically normalizing named entities, 2) organizing Statements by specificity, and 3) exploiting information about Statement sources, frequency and specificity to predict Statement reliability. Others have introduced innovative methods for using machine reading and curated databases for automated model learning and extension, while also making use of INDRA to process reader output (Holtzapple et al., 2020) and estimate Statement reliability (Ahmed et al., 2021). We believe our work to be the first demonstration of a method that automatically assembles reliable, non-redundant mechanistic knowledge from both curated resources and multiple biomedical text mining systems.

Our approach focuses on capturing the types of information typically represented in biological pathway databases: post-translational modifications and physical and regulatory interactions among proteins, chemicals, and biological processes. It does not currently represent genetic interactions, gene-disease relationships, biomarkers, or other types of statistical associations. However, given suitable data sources and text extraction systems, the same approaches for named entity linking, hierarchical assembly and reliability assessment can be used for these types of information as well. Indeed, the core methodology described here has been used to generate probabilistic causal models from a reading system that extracts causal relations from open-domain text (Sharp et al., 2019).

Automatically assembled knowledge bases have many uses in computational biology beyond the biocuration and functional genomics use cases we described here. A number of methods have been described that use pathway information for regularization in machine learning (Sokolov et al., 2016), to control false discovery in hypothesis testing (Babur et al., 2015), and to generate causal hypotheses from -omics data (Dugourd et al., 2021; Tuncbag et al., 2016). Most current methods for prior-guided data analysis require information about mechanisms to be “flattened” into a directed (possibly signed) networks (as we did in for our gene dependency correlation analysis). However, INDRA offers the ability to assemble information from multiple sources while preserving much of the information about mutations, modifications, and activity states that are necessary for detailed modeling. This supports the further development of analytical methods that exploit prior knowledge that is both broad and mechanistically detailed. INDRA facilitates this because it assembles information from sources in terms of knowledge-level assertions rather than model-specific implementations, different types of causal models can be generated from the assembled knowledge depending on the downstream application, including signed directed graphs, Boolean networks, or other types of executable models. In our previous work, we described a method for automatically assembling curated natural language text into detailed dynamical signaling models (Gyori et al., 2017). In principle, the methods described here allow for mechanistically detailed signaling models to be initialized from systematically compiled knowledge bases, with a quality suitable for static causal analysis ((Gyori et al., 2021), see https://emmaa.indra.bio). However, manual curation is generally still required to produce dynamical simulation models from automatically assembled assertions, due to the need to supply reverse rates and guarantee detailed balance; making this process more efficient is an area of ongoing research (Gyori and Bachman, 2021).

One of the striking conclusions from this work is that different reading systems extract very different types of information from the same text corpus. Moreover, even in the case of a single INDRA Statement, different reading systems extract different mentions from the same text. This points to the value of using multiple readers in parallel, something that has not previously been widely explored and suggests that direct comparison of reading system errors has the potential to improve these systems. To make use of multiple readers we developed an approach for estimating the technical reliability of Statements based on the number of mentions, the characteristics of their supporting evidences, and the properties of individual reading systems. While addressing this purely technical source of uncertainty is a prerequisite for the practical use of text mined information in downstream applications, addressing additional types of uncertainty in assembled knowledge and models is a worthwhile area of future research. In particular, there is a need for systematic approaches to managing conflicts and contradictions among assembled Statements (**Figure 4A**, upper right), which often take the form of polarity conflicts (“A activates B” vs. “A inhibits B”). While polarity conflicts can arise due to systematic errors in machine reading (Noriega-Atala et al., 2019), many represent inconsistent reports from the underlying scientific literature. These conflicts can potentially be addressed by a more thorough incorporation of biological context alongside causal information (Noriega-Atala et al., 2020), through the use of functional data such as the DepMap, or potentially by ensemble modeling procedures that capture polarity uncertainty in downstream analysis. Another primary concern in the use of text-mined information is the unreliability of many scientific studies (Baker, 2016). Recent efforts in meta-scientific analysis have examined features such as journal impact factor, article citations, and collaboration networks among researchers to determine whether these can predict the likelihood of the future replication of a study (Danchev et al., 2019). Large-scale assembly of causal information from the literature has the potential to facilitate the study of both biological and meta-scientific sources of scientific contradictions.

It is interesting to speculate what might be possible were all of PubMed to be made fully machine readable. The corpus of 570,000 papers used in this study were chosen in part because they focus on human genes and their functions. Because it is not a randomly selected subset of all 30 million PubMed articles, comprehensive machine reading followed by assembly in INDRA is unlikely to generate 60-fold more mechanistic information than the current study. To obtain a rough estimate of what could be expected, we determined the increase in the number of unique Statements and total mentions for a single gene of interest, BRAF, obtainable by processing all machine-readable abstracts and full text articles in PubMed with two readers, Reach and Sparser. We found that, relative to the Benchmark Corpus, unique Statements roughly doubled (from ∼1,500 to ∼3,300), while total mentions tripled (∼4,000 to ∼12,000) and the total number of supporting articles quadrupled (∼1,000 to ∼4,000). These numbers highlight the potential value of applying knowledge extraction and assembly methods more broadly. However, in performing this analysis we were limited by the availability of full text content, as we were in our assembly of the Benchmark Corpus (Table 1). The contribution of text mining tools to machine readable knowledge is expected to be much more significant when a greater proportion of full text scientific articles are both legally and technically accessible (Westergaard et al., 2018).

## ACKNOWLEDGEMENTS

This work was supported by DARPA grants W911NF-15-1-0544 and W911NF-20-1-0255 and by NCI grant U54-CA225088. We are grateful to the developers of Reach, Sparser, MedScan, RLIMS-P, TRIPS, and ISI/AMR for their support in integrating their reading systems; to members of the Laboratory of Systems Pharmacology Klas Karis, Albert Steppi, Charles Hoyt, Patrick Greene, Diana Kolusheva, and Daniel Milstein for their contributions to INDRA; and to Juliann Tefft for her helpful comments on the manuscript.

## AUTHOR CONTRIBUTIONS

JAB and BMG conceived and implemented the software, and performed the analysis. JAB, BMG and PKS wrote the paper.

## METHODS

### Article corpus for event extraction

The Entrez gene database was queried with the official gene symbols for all human genes in the HUGO database for MEDLINE articles curated as having relevance to the function of each gene. The resulting list of PubMed identifiers (PMIDs) is included in the code and data associated with the paper at https://github.com/sorgerlab/indra_assembly_paper. For these PMIDs, we obtained full text content when available from three sources: The PubMed Central open access and author’s manuscript collections, and the Elsevier Text and Data mining API (https://dev.elsevier.com). For the remaining PMIDs, we obtained abstracts from PubMed. Table 1 shows the distribution of text content sources.

### Text mining of article corpus

We used multiple text mining systems integrated with INDRA to process all or part of the corpus of interest described in the previous section.

#### Reach

version 1.3.3 was downloaded from https://github.com/clulab/reach and used to process all text content for the collected corpus described in the previous section. Reach reading output was processed into INDRA Statements using the indra.sources.reach module.

#### Sparser

was obtained as an executable image from its developers and was used to process all text content for the collected corpus described in the previous section. The Sparser source code is available at https://github.com/ddmcdonald/sparser and the Sparser executable is available as part of the INDRA Docker image which can be obtained from https://hub.docker.com/r/labsyspharm/indra. Sparser reading output was processed into INDRA Statements using the indra.sources.sparser module.

#### MedScan

reader output for the collected corpus described in the previous section was obtained from Elsevier and processed into INDRA Statements using the indra.sources.medscan module.

#### TRIPS/DRUM

was obtained from https://github.com/wdebeaum/drum and used to process part of the text content for the collected corpus, as follows. First, we selected all papers for which only an abstract was available, then selected those papers from which Reach, Sparser and MedScan extracted at least one Statement about any of 227 genes relevant for a key cancer signaling pathway, the Ras pathway. This resulted in a total of 42,158 abstracts which were processed with TRIPS/DRUM. The outputs were then processed into INDRA Statements using the indra.sources.trips module.

#### RLIMS-P

reader output for PubMed abstracts and PubMedCentral full text articles was obtained from https://hershey.dbi.udel.edu/textmining/export/ (accessed June 2019), and then filtered to the corpus of interest described in the previous section. The outputs were then processed into INDRA Statements using the indra.sources.rlimsp module.

#### ISI/AMR

(Docker image available at https://hub.docker.com/r/sahilgar/bigmechisi) reader output was provided by the system’s creators for 10,433 articles which were filtered to the corpus of interest resulting in a total of 1,878 reader outputs. These were then processed into INDRA Statements using the indra.sources.isi module.

### Structured sources

In addition to text mining, we processed multiple pathway databases with INDRA to obtain INDRA Statements.

#### TRRUST

release 4/16/2018 with human transcription factor-target relationships was obtained from https://www.grnpedia.org/trrust/data/trrust_rawdata.human.tsv and processed into INDRA Statements using the indra.sources.trrust module.

#### Signor

content was processed through the Signor web service (https://signor.uniroma2.it/download_entity.php) in June 2019 and processed into INDRA Statements using the indra.sources.signor module.

#### HPRD

content was obtained from http://www.hprd.org/RELEASE9/HPRD_FLAT_FILES_041310.tar.gz and processed into INDRA Statements using the indra.sources.hprd module.

#### BEL

content was obtained from the Selventa Large Corpus available at https://raw.githubusercontent.com/cthoyt/selventa-knowledge/master/selventa_knowledge/large_corpus.bel and processed using PyBEL and the indra.sources.bel module into INDRA Statements.

#### CausalBioNet

content was processed from JGF files from http://causalbionet.com/Content/jgf_bulk_files/Human-2.0.zip and processed into INDRA Statements using PyBEL and the indra.sources.bel module.

#### BioGRID

content was obtained from https://downloads.thebiogrid.org/Download/BioGRID/Release-Archive/BIOGRID-4.2.192/BIOGRID-ALL-4.2.192.tab3.zip and processed into INDRA Statements using the indra.sources.biogrid module.

#### PhosphoSitePlus

content was downloaded from https://www.phosphosite.org/staticDownloads in June 2019 via the “BioPAX:Kinase-substrate information” link, in BioPAX format, and processed into INDRA Statements using the indra.sourecs.biopax module.

#### Pathway Commons

content was obtained from https://www.pathwaycommons.org/archives/PC2/v12/PathwayCommons12.Detailed.BIOPAX.owl.gz and processed using PyBioPAX and the indra.sources.biopax module into INDRA Statements. To account for the fact that BioGRID, PhosphoSitePlus and HPRD content were obtained separately (and these are also available as part of Pathway Commons), we filtered out interactions from these sources when processing Pathway Commons.

The scripts to process each source as described above is available at: https://github.com/sorgerlab/indra_assembly_paper/blob/master/run_assembly/process_sources.py.

### Procedure for identifying duplicates and refinements

The INDRA ontology graph combines entries across multiple ontologies and represents each entry as a graph node with a set of properties (namespace, identifier, standard name). There are three types of edges in the graph: *xref* (cross-reference meaning that the source node and the target node, often from different namespaces, are equivalent), *isa* (the source node is one of a set of entities represented by the parent node), and *partof* (the source node is part of a complex represented by the parent node). Each INDRA Agent has zero or more namespace/identifier pairs associated with it which constitute its grounding.

When standardizing the grounding of INDRA Agents, the *xref* edges of the ontology graph are traversed following all directed paths starting from each available grounding for the Agent. The namespaces and identifiers of nodes visited along these paths are then added as grounding for the Agent. We then use a priority order of namespaces to assign a single *canonical grounding* to an Agent. If an Agent has no groundings available, its name is used as canonical grounding.

When determining whether two Statements are duplicates, we require that (1) the two Statements’ types are the same (2) all the Agent arguments of the two Statements are matching in their canonical grounding, (3) all states (activity, modifications, bound conditions, location, mutations) of the matching Agents of the two Statements are equivalent, and (4) all additional Statement arguments are equivalent (e.g., residue and position for a Modification Statement). To avoid making pairwise comparisons, we construct an equivalence key from properties (1-4) needed to determine equivalence for each individual Statement, and then use a hash map data structure to group Statements efficiently by equivalence key. Groups of Statements having the same equivalence key are collapsed into a single Statement and their Evidences are concatenated.

For finding refinements among Statements, we make use of the INDRA ontology graph’s *isa* and *partof* edges. For determining a refinement, we require that the two Statements have the same type, and that one Statement is a refinement of the other with respect to at least one of the properties (2-4) described above, and that the other Statement does not refine the first one based on any of these properties. In other words, if one Statement is more specific than the other according to one property but less specific according to another property, there is no refinement relationship between them at the Statement level.

### Binomial, beta-binomial, and INDRA Belief models of Statement reliability

#### The INDRA Belief Model

The “INDRA belief model” represents the probability of a Statement being correct as the result of a two step-random process (Figure 4C). The first process considers the probability that a Statement is drawn from the pool of Statements that are *always* incorrect, regardless of the number of evidences they have. This probability is based on the *systematic error* parameter for each reading system. If the Statement is *not* from this pool, then its reliability is alternatively modeled to follow a binomial distribution assuming a particular *random error* rate for that source. Like the beta-binomial model, the INDRA belief model captures the plateau in Statement reliability (Figure 4D), though the predicted distributions for evidence correctness do not correspond well to the empirical U-shaped distribution (Figure S4A).

The INDRA Belief Model is calculated based on evidences belonging to a Statement, each evidence having been produced by a source such as a text mining system or a pathway database integrated with INDRA. In the simple case of a single knowledge source, we define the belief of a

Statement as

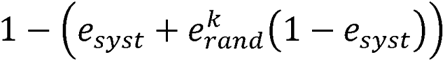

where *e_syst_* and *e_rand_* are the systematic and random error parameters for the given source, respectively. This model can also be generalized to multiple sources as follows. Assume there are a total of *K* known sources *S =* {*S_1_*, *S_2_*, …, *S_K_*} each associated with a random and systematic error rate. For source *S_k_ E_k,syst_* will denote the systematic error rate, and *e_k,rand_* the random error rate. Given a set of *N* evidences E*T =* {*E_T_*_,1_, …, *E_T,N_*}, with Source (*E_T,i_*) ∈ *S* corresponding to the source of evidence of *E_T,n_*, we introduce *N_T,k_*, the number of evidences for Statement T from source *S_k:_*

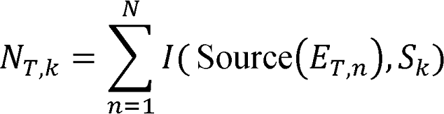

where *I*(*X*, *Y*) stands for the indicator function which evaluates to 1 if *X* = *Y*, and 0 otherwise. We then define the belief of Statement T as follows:

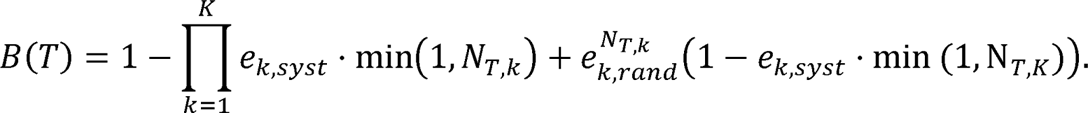

For the calculation of beliefs for a Statement that is refined by other Statements, we introduce the extended evidence set denoted as *E’*(*T*)which is defined as

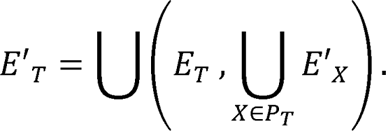

Here *X* ∈ *P_T_* if and only if *X*refines *T*. In other words, we take the union of all pieces of evidence for the Statement itself and all the Statements by which it is refined, recursively. We then apply the equation for *N_T,k_* and *B*(*T*) to *E’*(*T*) instead of *E’*(*T*) in the obvious way.

When the quality of fit of the three different models was compared using maximum likelihood parameter values, the original belief model performed very slightly better than the beta-binomial model for both the Reach and Sparser reading systems (Table 3).

#### The Binomial and Beta-binomial belief models

The binomial model treats every individual evidence sentence as a Bernoulli trial, where the probability of a single reading system being jointly incorrect for all sentences decreases according to a binomial distribution (e.g., the probability of incorrectly processing ten sentences is analogous to flipping a coin ten times and getting ten tails). The binomial model substantially overestimates the reliability of Statements with three or more evidences from Reach, due to the fact that it does not account for systematic errors (Figure 4A). In addition, the binomial model predicts that for a Statement with *k* evidences, the mode of the distribution of number of correct evidences is close to *k/2* (bell shaped red curves in Figure S4B), whereas the curation data shows that evidences are more likely to be either all incorrect (zero bars) or all correct (right-most bars).

The binomial belief model has a single random error rate parameter *e_rand_* for each source, and –making use of definitions from the previous section – the belief for a Statement with evidences from multiple sources can be calculated as

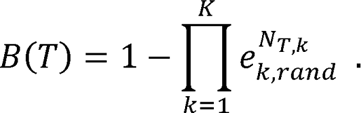

The beta-binomial model is based on a binomial model where the probability of each mention being correctly extracted is itself drawn from a beta distribution (Wilcox, 1979). The beta-binomial model better captures the tendency of Statement reliability to plateau below 100% (Figure 4D) as well as the U-shaped distributions of the numbers of underlying correct evidences (Figure S4C).

The beta-binomial belief model has two parameters for each source, *α* and *β*, and for a Statement with evidences from multiple sources, it can be calculated as

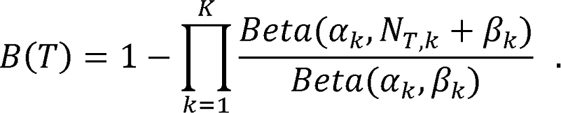

where *Beta* is the standard beta-function.

### Machine-learned models of Statement reliability

#### Model types and evaluation

Classification models evaluated for their ability to predict Statement correctness were obtained from the Python package *sklearn.* Evaluated models included Support Vector Classification (*sklearn.svm.SVC* with probability estimation enabled), k-Nearest Neighbors (*sklearn.neighbors.KNeighborsClassifier,* used with default parameters), logistic regression with log-transformed mention counts (*sklearn.linear_model.LogisticRegression)*, and Random Forests (*sklearn.ensemble.RandomForestClassifier* with *n_estimators=2000* and *max_depth=13,* obtained by manual hyper-parameter optimization). Model performance was evaluated by 10-fold cross-validation; each fold was used to calculate the area under the precision-recall curve (AUPRC) for the held-out data. Values in Table 3 reflect the mean and standard deviations of AUPRC values across the 10 folds.

#### Encoding of features for Statement belief prediction

##### Reader mention counts

Mention counts for each reader were included as distinct features (columns) for each Statement. When incorporating evidence from more specific Statements (“specific evidences” in Table 3) these were added in a separate set of columns for each reader; a Statement could thus have two columns with Reach mention counts, one for mentions directly supporting the Statement, and another for mentions obtained from more specific Statements.

###### Number of unique PMIDs

Unique PMIDs supporting each Statement were obtained from its mentions and added as a single feature.

###### Statement type

Statement types were one-hot encoded (one binary feature for each type, Activation, Inhibition, Phosphorylation, etc.)

###### Average evidence length

Mention texts directly supporting the Statement were split by whitespace; the number of resulting substrings were counted and averaged across all mentions and included as a feature. *“Promoter” frequency.* The number of mention texts containing the term “promoter” were counted and the resulting value was divided by the total number of mentions to obtain a frequency of the occurrence of this keyword.

### Availability of data and material

The datasets generated and analyzed during the current study, as well as the source code used to generate results is available in the repository https://github.com/sorgerlab/indra_assembly_paper. INDRA is available at https://github.com/sorgerlab/indra under an open-source BSD 2-clause license.

---

**Figure S1.**
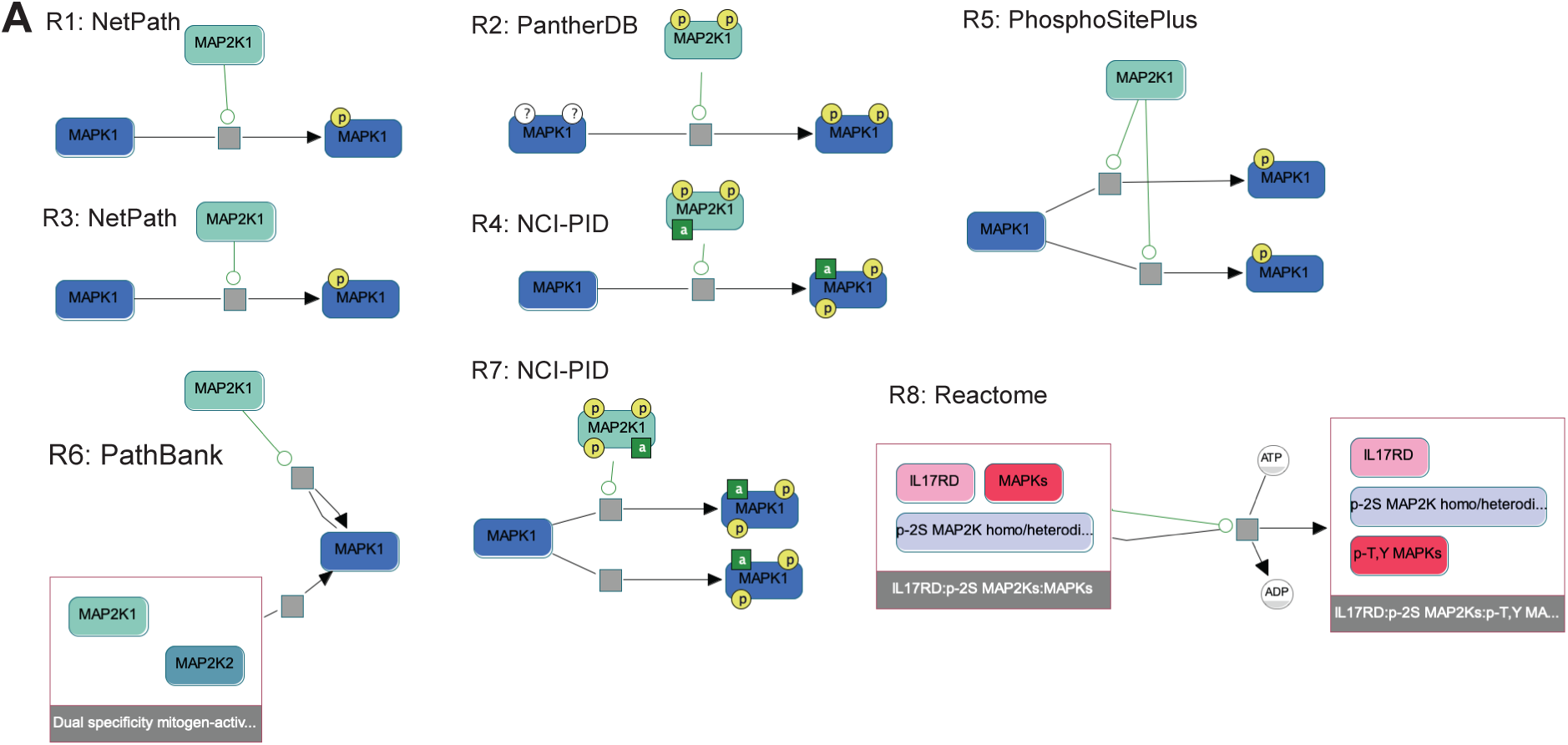
Differences in curation practices across databases integrated by Pathway Commons. (A) A subgraph of the “paths-from-to” query between MAP2K1 and MAPK1 obtained from Pathway Commons and visualized using the ChiBE software (Babur et al., 2009). Each biochemical reaction (R1-R8) depicts a different curation of the same reaction in which MAP2K1 phosphorylates MAPK1. The original source database (e.g., NetPath) is shown next to each reaction. Inconsistencies include (i) the reaction structure itself, with some specifying a single step phosphorylation of two sites (e.g., R4) while others specify single-site phosphorylation (e.g., R1), or the explicit representation of ADP and ATP as part of the reaction (R8); (ii) the phosphorylation status of MAP2K1, with no phosphorylation status given in R1, R3, R5, R6 and R8, two phosphorylation sites indicated in R2, R4, and three phosphorylation sites in R7; (iii) the initial state of MAPK1, with R2 explicitly indicating unphosphorylated status, while other reactions do not make this explicit; (iv) the final state of MAPK1, with some reactions representing MAPK1 phosphorylation on an unspecified site (R1, R3), and others providing specific phosphorylation sites (e.g., R2); (v) the specification of active states, with R4 being the only reaction representing MAP2K1 explicitly as active, while R4 and R7 are the only reactions specifying that MAPK1 is active after phosphorylation; (vi) the presence of other co-factors such as IL17RD (R8) as part of the reaction.

**Figure S2.**
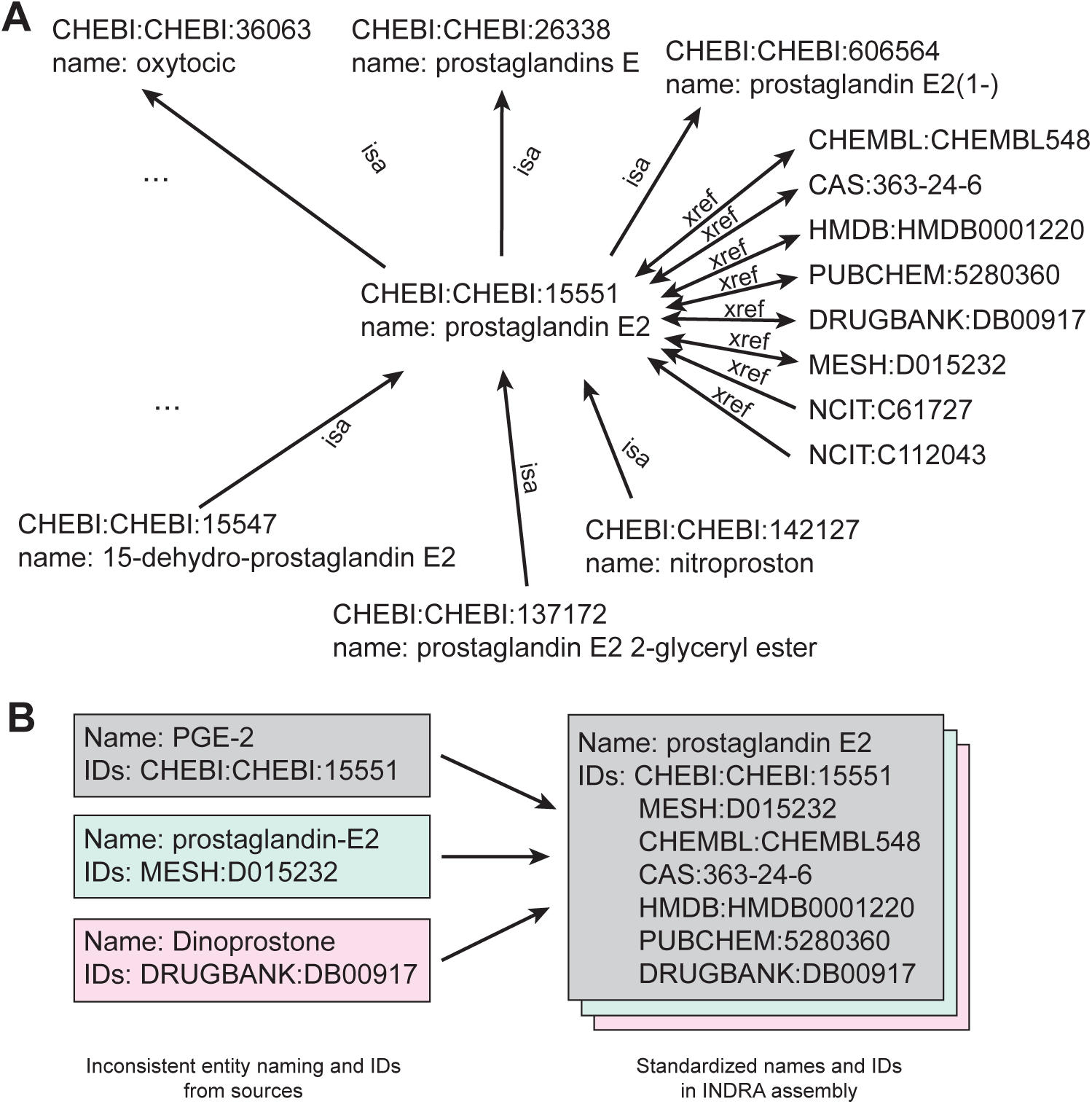
Ontology graph guiding INDRA knowledge assembly. (A) A subgraph of the INDRA ontology graph showing the neighborhood of the node representing “prostaglandin E2” in the ChEBI database (CHEBI:15551). Edges represent “isa” relationships to more general terms (and from more specific terms), and “xref” edges represent identifier equivalence to nodes representing entries in other databases including MeSH, DrugBank, ChEMBL, CAS, PubChem, and NCIT. Each ontology graph node also provides a name that can be used for standardization and display purposes. (B) Example of three entities with inconsistent names and identifiers which, when standardized by INDRA using the ontology graph, are normalized to consistent entities with identical names and sets of identifiers.

**Figure S4.**
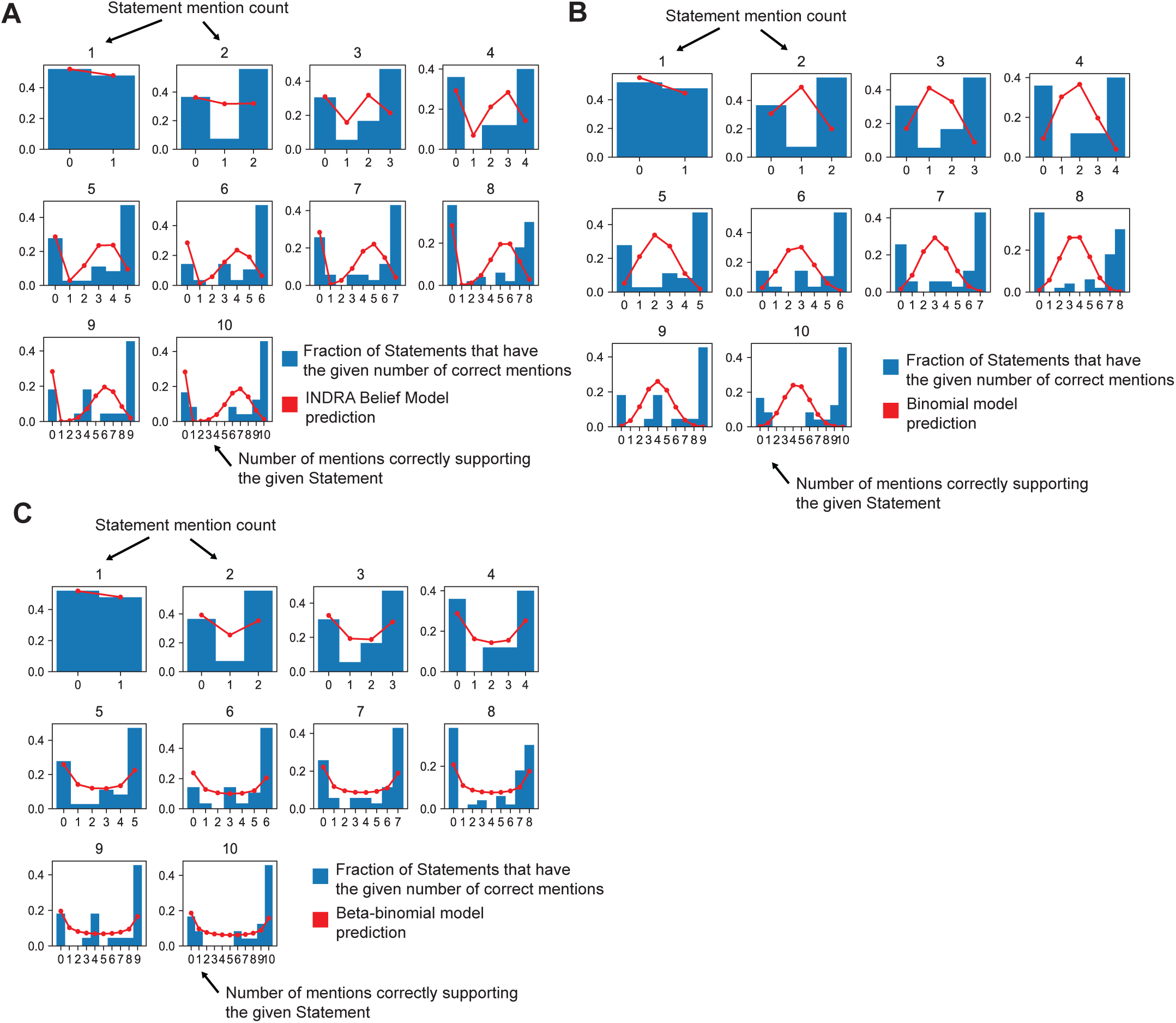
Observed and predicted distributions of mentions correctly extracted by Reach for Statements supported by up to 10 Reach mentions. (A) Frequencies of correct mentions predicted by the INDRA Belief Model. The blue bars in each subplot show the frequencies of statements with *k* correctly extracted mentions for *n* total mentions for the Statement (considering mentions from the Reach reader only). The red line in each subplot shows the frequencies of correct mentions expected by the INDRA Belief Model. The INDRA Belief Model expects a substantial proportion of Statements to have an intermediate number of correctly extracted mentions, whereas the empirical data suggests that Statements are more likely to be associated with mentions that are either all correct or incorrect. (B). Frequencies of correct mentions expected by the Binomial model. Blue bars are identical to (A). (C). Frequencies of correct mentions expected by the Beta-binomial model. Blue bars are identical to (A) and (B). The Beta-binomial model differs from the INDRA Belief Model and Binomial models in that it predicts relatively greater proportions of Statements with mentions that are either all correct or incorrect.

**Figure S5.**
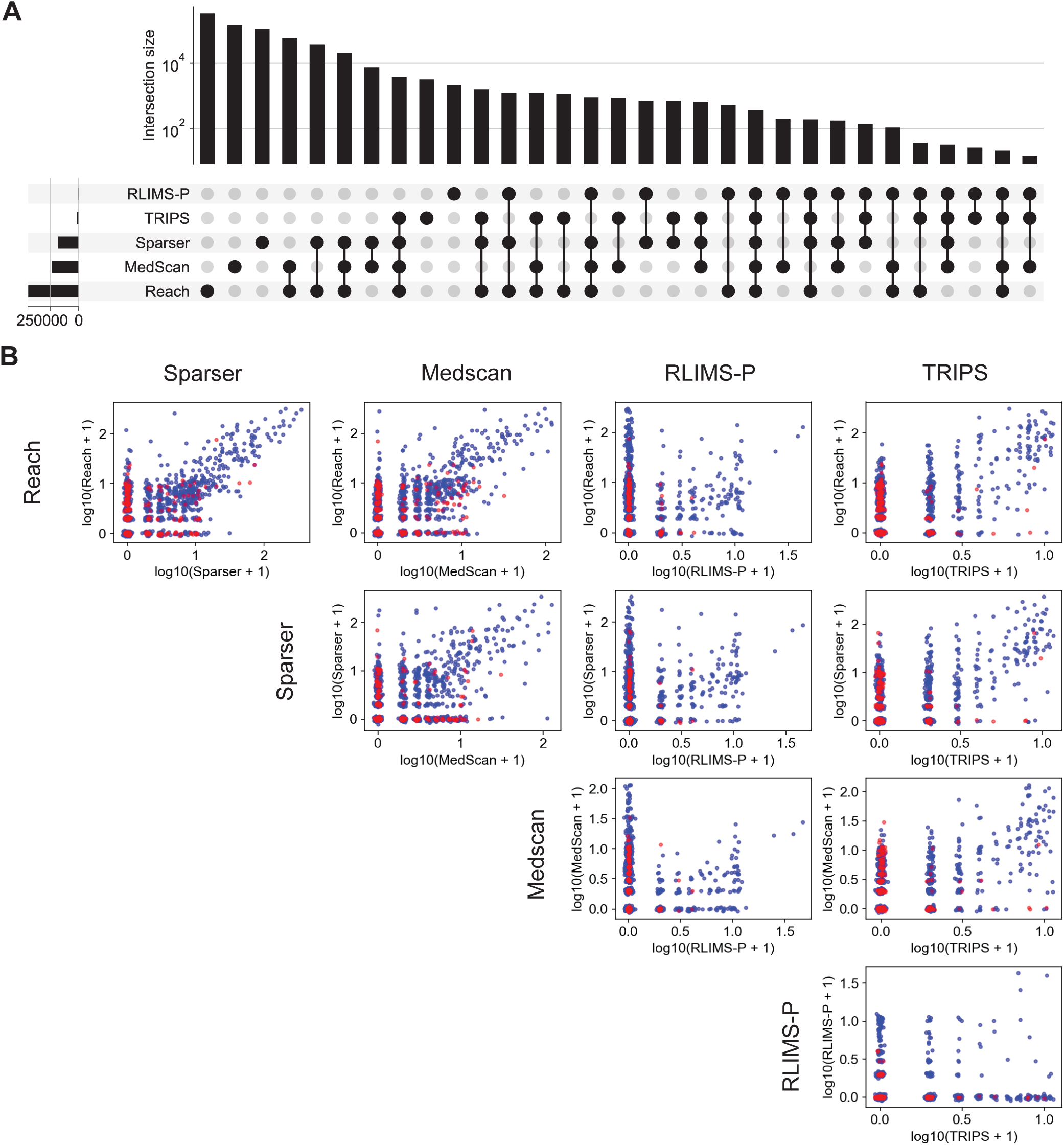
Reader overlap and Statement correctness. (A) Upset plot (equivalent to a Venn diagram with more than 3 sets) of Statement support for five machine reading systems integrated by INDRA. Data is identical to Figure 5A but intersection sizes are plotted on a log scale and all 32 possible reader combinations are shown. (B) Multi-reader mention counts and Statement correctness. Each subplot shows the relationship between mention counts from a combination of two readers for manually curated Statements. Blue points represent Statements that were curated as correct; red points were curated as incorrect. A small amount of random jitter has been added to each point to indicate the density of points with fewer mention counts.

## REFERENCES

Ahmed, Y., Telmer, C.A., and Miskov-Zivanov, N. (2021). CLARINET: Efficient learning of dynamic network models from literature. Bioinforma. Adv. 1, vbab006.

Allen, J., de Beaumont, W., Galescu, L., and Teng, C.M. (2015). Complex Event Extraction using DRUM. ACL-IJCNLP 2015 1–11..

Alstott, J., Bullmore, E., and Plenz, D. (2014). powerlaw: A Python Package for Analysis of Heavy-Tailed Distributions. PLoS ONE 9, e85777. https://doi.org/10.1371/journal.pone.0085777.

Ananiadou, S., Thompson, P., Nawaz, R., McNaught, J., and Kell, D.B. (2015). Event-based text mining for biology and functional genomics. Brief. Funct. Genomics 14, 213–230. https://doi.org/10.1093/bfgp/elu015.

Ashburner, M., Ball, C.A., Blake, J.A., Botstein, D., Butler, H., Cherry, J.M., Davis, A.P., Dolinski, K., Dwight, S.S., Eppig, J.T., et al. (2000). Gene Ontology: Tool for the Unification of Biology. The Gene Ontology Consortium. Nat. Genet. 25, 25–29. https://doi.org/10.1038/75556.

Babur, O., Dogrusoz, U., Demir, E., and Sander, C. (2009). ChiBE: interactive visualization and manipulation of BioPAX pathway models. Bioinformatics 26, 429–431..

Babur, Ö., Gönen, M., Aksoy, B.A., Schultz, N., Ciriello, G., Sander, C., and Demir, E. (2015). Systematic identification of cancer driving signaling pathways based on mutual exclusivity of genomic alterations. Genome Biol. 16, 45. https://doi.org/10.1186/s13059-015-0612-6.

Babur, Ö., Luna, A., Korkut, A., Durupinar, F., Siper, M.C., Dogrusoz, U., Vaca Jacome, A.S., Peckner, R., Christianson, K.E., Jaffe, J.D., et al. (2021). Causal interactions from proteomic profiles: Molecular data meet pathway knowledge. Patterns N. Y. N 2, 100257. https://doi.org/10.1016/j.patter.2021.100257.

Bachman, J.A., Gyori, B.M., and Sorger, P.K. (2018). FamPlex: A Resource for Entity Recognition and Relationship Resolution of Human Protein Families and Complexes in Biomedical Text Mining. BMC Bioinformatics 19, 248. https://doi.org/10.1186/s12859-018-2211-5.

Bachman, J.A., Gyori, B.M., and Sorger, P.K. (2019). Assembling a phosphoproteomic knowledge base using ProtMapper to normalize phosphosite information from databases and text mining. BioRxiv 822668. https://doi.org/10.1101/822668.

Baker, M. (2016). 1,500 scientists lift the lid on reproducibility. Nature 533, 452–454. https://doi.org/10.1038/533452a.

Benseddik, K., Sen Nkwe, N., Daou, P., Verdier-Pinard, P., and Badache, A. (2013). ErbB2-dependent chemotaxis requires microtubule capture and stabilization coordinated by distinct signaling pathways. PloS One 8, e55211. https://doi.org/10.1371/journal.pone.0055211.

Bourne, P.E., Lorsch, J.R., and Green, E.D. (2015). Perspective: Sustaining the Big-Data Ecosystem. Nature 527, S16–17. https://doi.org/10.1038/527S16a.

Cerami, E.G., Gross, B.E., Demir, E., Rodchenkov, I., Babur, O., Anwar, N., Schultz, N., Bader, G.D., and Sander, C. (2011). Pathway Commons, a web resource for biological pathway data. Nucleic Acids Res. 39, D685–690. https://doi.org/10.1093/nar/gkq1039.

Clauset, A., Shalizi, C.R., and Newman, M.E.J. (2009). Power-Law Distributions in Empirical Data. SIAM Rev. 51, 661–703. https://doi.org/10.1137/070710111.

Coppari, E., Yamada, T., Bizzarri, A.R., Beattie, C.W., and Cannistraro, S. (2014). A nanotechnological, molecular-modeling, and immunological approach to study the interaction of the anti-tumorigenic peptide p28 with the p53 family of proteins. Int. J. Nanomedicine 9, 1799–1813. https://doi.org/10.2147/IJN.S58465.

Craver, C.F., and Darden, L. (2013). In Search of Mechanisms: Discoveries across the Life Sciences (University of Chicago Press).

Danchev, V., Rzhetsky, A., and Evans, J.A. (2019). Centralized scientific communities are less likely to generate replicable results. ELife 8, e43094. https://doi.org/10.7554/eLife.43094.

Demir, E., Cary, M.P., Paley, S., Fukuda, K., Lemer, C., Vastrik, I., Wu, G., D’Eustachio, P., Schaefer, C., Luciano, J., et al. (2010). BioPAX – A Community Standard for Pathway Data Sharing. Nat. Biotechnol. 28, 935– 942. https://doi.org/10.1038/nbt.1666.

Dempster, J.M., Pacini, C., Pantel, S., Behan, F.M., Green, T., Krill-Burger, J., Beaver, C.M., Younger, S.T., Zhivich, V., Najgebauer, H., et al. (2019). Agreement between two large pan-cancer CRISPR-Cas9 gene dependency data sets. Nat. Commun. 10, 5817. https://doi.org/10.1038/s41467-019-13805-y.

Deng, Y., Bhattacharya, S., Swamy, O.R., Tandon, R., Wang, Y., Janda, R., and Riedel, H. (2003). Growth factor receptor-binding protein 10 (Grb10) as a partner of phosphatidylinositol 3-kinase in metabolic insulin action. J. Biol. Chem. 278, 39311–39322. https://doi.org/10.1074/jbc.M304599200.

Doherty, L.M., Mills, C.E., Boswell, S.A., Liu, X., Hoyt, C.T., Gyori, B.M., Buhrlage, S.J., and Sorger, P.K. (2021). Integrating multi-omics data reveals function and therapeutic potential of deubiquitinating enzymes. https://doi.org/10.1101/2021.08.06.455458.

Dugourd, A., Kuppe, C., Sciacovelli, M., Gjerga, E., Gabor, A., Emdal, K.B., Vieira, V., Bekker-Jensen, D.B., Kranz, J., Bindels, E.M.J., et al. (2021). Causal integration of multi-omics data with prior knowledge to generate mechanistic hypotheses. Mol. Syst. Biol. 17, e9730. https://doi.org/10.15252/msb.20209730.

Fabregat, A., Jupe, S., Matthews, L., Sidiropoulos, K., Gillespie, M., Garapati, P., Haw, R., Jassal, B., Korninger, F., May, B., et al. (2018). The Reactome Pathway Knowledgebase. Nucleic Acids Res. 46, D649–D655. https://doi.org/10.1093/nar/gkx1132.

Foreman-Mackey, D., Hogg, D.W., Lang, D., and Goodman, J. (2013). emcee: The MCMC Hammer. Publ. Astron. Soc. Pac. 125, 306–312. https://doi.org/10.1086/670067.

Forscher, B.K. (1963). Chaos in the Brickyard. Science 142, 339–339. https://doi.org/10.1126/science.142.3590.339.

Frank, S.R., Köllmann, C.P., van Lidth de Jeude, J.F., Thiagarajah, J.R., Engelholm, L.H., Frödin, M., and Hansen, S.H. (2017). The focal adhesion-associated proteins DOCK5 and GIT2 comprise a rheostat in control of epithelial invasion. Oncogene 36, 1816–1828. https://doi.org/10.1038/onc.2016.345.

Garg, S., Galstyan, A., Hermjakob, U., and Marcu, D. (2016). Extracting biomolecular interactions using semantic parsing of biomedical text. In Thirtieth AAAI Conference on Artificial Intelligence, (Phoenix, Arizona), pp. 2718–2726.

Glavaški, M., and Velicki, L. (2021). Humans and machines in biomedical knowledge curation: hypertrophic cardiomyopathy molecular mechanisms’ representation. BioData Min. 14, 45. https://doi.org/10.1186/s13040-021-00279-2.

Gyori, B.M., and Bachman, J.A. (2021). From knowledge to models: Automated modeling in systems and synthetic biology. Curr. Opin. Syst. Biol. 100362. https://doi.org/10.1016/j.coisb.2021.100362.

Gyori, B.M., Bachman, J.A., Subramanian, K., Muhlich, J.L., Galescu, L., and Sorger, P.K. (2017). From word models to executable models of signaling networks using automated assembly. Mol. Syst. Biol. 13, 954..

Gyori, B.M., Bachman, J.A., and Kolusheva, D. (2021). A self-updating causal model of COVID-19 mechanisms built from the scientific literature. In BioCreative VII Challenge Evaluation Workshop, p. 249.

Gyori, B.M., Hoyt, C.T., and Steppi, A. (2022). Gilda: biomedical entity text normalization with machine-learned disambiguation as a service. Bioinforma. Adv. 2, vbac034. https://doi.org/10.1093/bioadv/vbac034.

Hastings, J., Owen, G., Dekker, A., Ennis, M., Kale, N., Muthukrishnan, V., Turner, S., Swainston, N., Mendes, P., and Steinbeck, C. (2016). ChEBI in 2016: Improved services and an expanding collection of metabolites. Nucleic Acids Res. 44, D1214–D1219. https://doi.org/10.1093/nar/gkv1031.

Himmelstein, D.S., Lizee, A., Hessler, C., Brueggeman, L., Chen, S.L., Hadley, D., Green, A., Khankhanian, P., and Baranzini, S.E. (2017). Systematic integration of biomedical knowledge prioritizes drugs for repurposing. ELife 6, e26726. https://doi.org/10.7554/eLife.26726.

Holtzapple, E., Telmer, C.A., and Miskov-Zivanov, N. (2020). FLUTE: Fast and reliable knowledge retrieval from biomedical literature. Database J. Biol. Databases Curation 2020. https://doi.org/10.1093/database/baaa056.

Hoyt, C.T., Domingo-Fernández, D., Aldisi, R., Xu, L., Kolpeja, K., Spalek, S., Wollert, E., Bachman, J., Gyori, B.M., Greene, P., et al. (2019a). Re-curation and rational enrichment of knowledge graphs in Biological Expression Language. Database 2019, baz068. https://doi.org/10.1093/database/baz068.

Hoyt, C.T., Domingo-Fernández, D., Mubeen, S., Llaó, J.M., Konotopez, A., Ebeling, C., Birkenbihl, C., Muslu, Ö., English, B., Müller, S., et al. (2019b). Integration of Structured Biological Data Sources using Biological Expression Language. BioRxiv 631812. https://doi.org/10.1101/631812.

Hucka, M., Finney, A., Sauro, H.M., Bolouri, H., Doyle, J.C., Kitano, H., Arkin, A.P., Bornstein, B.J., Bray, D., Cornish-Bowden, A., et al. (2003). The Systems Biology Markup Language (SBML): A Medium for Representation and Exchange of Biochemical Network Models. Bioinformatics 19, 524–531. https://doi.org/10.1093/bioinformatics/btg015.

Ietswaart, R., Gyori, B.M., Bachman, J.A., Sorger, P.K., and Churchman, L.S. (2021). GeneWalk identifies relevant gene functions for a biological context using network representation learning. Genome Biol. 22, 55. https://doi.org/10.1186/s13059-021-02264-8.

Islamaj Doğan, R., Kim, S., Chatr-aryamontri, A., Wei, C.-H., Comeau, D.C., Antunes, R., Matos, S., Chen, Q., Elangovan, A., Panyam, N.C., et al. (2019). Overview of the BioCreative VI Precision Medicine Track: mining protein interactions and mutations for precision medicine. Database 2019. https://doi.org/10.1093/database/bay147.

Jensen, L.J., Kuhn, M., Stark, M., Chaffron, S., Creevey, C., Muller, J., Doerks, T., Julien, P., Roth, A., Simonovic, M., et al. (2009). STRING 8–a Global View on Proteins and Their Functional Interactions in 630 Organisms. Nucleic Acids Res. 37, D412–416. https://doi.org/10.1093/nar/gkn760.

Jia, S., Xiang, Y., and Chen, X. (2019). Triple Trustworthiness Measurement for Knowledge Graph. World Wide Web Conf. - WWW 19 2865–2871. https://doi.org/10.1145/3308558.3313586.

Keegan, A.D., Zamorano, J., Keselman, A., and Heller, N.M. (2018). IL-4 and IL-13 Receptor Signaling From 4PS to Insulin Receptor Substrate 2: There and Back Again, a Historical View. Front. Immunol. 9, 1037. https://doi.org/10.3389/fimmu.2018.01037.

Kemper, B., Matsuzaki, T., Matsuoka, Y., Tsuruoka, Y., Kitano, H., Ananiadou, S., and Tsujii, J. ’ichi (2010). PathText: A Text Mining Integrator for Biological Pathway Visualizations. Bioinforma. Oxf. Engl. 26, i374–381. https://doi.org/10.1093/bioinformatics/btq221.

Kumar, S.M., Dai, J., Li, S., Yang, R., Yu, H., Nathanson, K.L., Liu, S., Zhou, H., Guo, J., and Xu, X. (2014). Human skin neural crest progenitor cells are susceptible to BRAFV600E-induced transformation. Oncogene 33, 832–841. https://doi.org/10.1038/onc.2012.642.

Lee, P.L., Ohlson, M.B., and Pfeffer, S.R. (2015). Rab6 regulation of the kinesin family KIF1C motor domain contributes to Golgi tethering. ELife 4. https://doi.org/10.7554/eLife.06029.

Lopez, C.F., Muhlich, J.L., Bachman, J.A., and Sorger, P.K. (2013). Programming biological models in Python using PySB. Mol. Syst. Biol. 9, 646. https://doi.org/10.1038/msb.2013.1.

Madan, S., Szostak, J., Komandur Elayavilli, R., Tsai, R.T.-H., Ali, M., Qian, L., Rastegar-Mojarad, M., Hoeng, J., and Fluck, J. (2019). The extraction of complex relationships and their conversion to biological expression language (BEL) overview of the BioCreative VI (2017) BEL track. Database J. Biol. Databases Curation 2019, baz084. https://doi.org/10.1093/database/baz084.

McDonald, D.D., Friedman, S.E., Paullada, A., Bobrow, R., and Burstein, M.H. (2016). Extending Biology Models with Deep NLP over Scientific Articles. In AAAI Workshop: Knowledge Extraction from Text, p.

Meyers, R.M., Bryan, J.G., McFarland, J.M., Weir, B.A., Sizemore, A.E., Xu, H., Dharia, N.V., Montgomery, P.G., Cowley, G.S., Pantel, S., et al. (2017). Computational correction of copy number effect improves specificity of CRISPR–Cas9 essentiality screens in cancer cells. Nat. Genet. 49, 1779–1784. https://doi.org/10.1038/ng.3984.

Mishra, G.R. (2006). Human protein reference database--2006 update. Nucleic Acids Res. 34, D411–D414. https://doi.org/10.1093/nar/gkj141.

Moody, T.W., Di Florio, A., and Jensen, R.T. (2012). PYK-2 is Tyrosine Phosphorylated after Activation of Pituitary Adenylate Cyclase Activating Polypeptide Receptors in Lung Cancer Cells. J. Mol. Neurosci. 48, 660– 666. https://doi.org/10.1007/s12031-012-9785-6.

Moret, N., Liu, C., Gyori, B.M., Bachman, J.A., Steppi, A., Hug, C., Taujale, R., Huang, L.-C., Berginski, M.E., Gomez, S.M., et al. (2021). A resource for exploring the understudied human kinome for research and therapeutic opportunities. BioRxiv https://doi.org/10.1101/2020.04.02.022277.

Mori, K., Giovannone, B., and Smith, R.J. (2005). Distinct Grb10 domain requirements for effects on glucose uptake and insulin signaling. Mol. Cell. Endocrinol. 230, 39–50. https://doi.org/10.1016/j.mce.2004.11.004.

Neil, D., Briody, J., Lacoste, A., Sim, A., Creed, P., and Saffari, A. (2018). Interpretable Graph Convolutional Neural Networks for Inference on Noisy Knowledge Graphs. ArXiv181200279 Cs Stat.

Noriega-Atala, E., Liang, Z., Bachman, J., Morrison, C., and Surdeanu, M. (2019). Understanding the Polarity of Events in the Biomedical Literature: Deep Learning vs. Linguistically-informed Methods. In Proceedings of the Workshop on Extracting Structured Knowledge from Scientific Publications, (Minneapolis, Minnesota: Association for Computational Linguistics), pp. 21–30.

Noriega-Atala, E., Hein, P.D., Thumsi, S.S., Wong, Z., Wang, X., Hendryx, S.M., and Morrison, C.T. (2020). Extracting Inter-Sentence Relations for Associating Biological Context with Events in Biomedical Texts. IEEE/ACM Trans. Comput. Biol. Bioinform. 17, 1895–1906. https://doi.org/10.1109/TCBB.2019.2904231.

Novichkova, S., Egorov, S., and Daraselia, N. (2003). MedScan, a Natural Language Processing Engine for MEDLINE Abstracts. Bioinforma. Oxf. Engl. 19, 1699–1706..

Nunes, K.P., Rigsby, C.S., and Webb, R.C. (2010). RhoA/Rho-kinase and vascular diseases: what is the link? Cell. Mol. Life Sci. CMLS 67, 3823–3836. https://doi.org/10.1007/s00018-010-0460-1.

Ostaszewski, M., Niarakis, A., Mazein, A., Kuperstein, I., Phair, R., Orta-Resendiz, A., Singh, V., Aghamiri, S.S., Acencio, M.L., Glaab, E., et al. (2021). COVID19 Disease Map, a computational knowledge repository of virus-host interaction mechanisms. Mol. Syst. Biol. 17, e10387. https://doi.org/10.15252/msb.202110387.

Oughtred, R., Stark, C., Breitkreutz, B.-J., Rust, J., Boucher, L., Chang, C., Kolas, N., O’Donnell, L., Leung, G., McAdam, R., et al. (2019). The BioGRID interaction database: 2019 update. Nucleic Acids Res. 47, D529–D541. https://doi.org/10.1093/nar/gky1079.

Pan, J., Meyers, R.M., Michel, B.C., Mashtalir, N., Sizemore, A.E., Wells, J.N., Cassel, S.H., Vazquez, F., Weir, B.A., Hahn, W.C., et al. (2018). Interrogation of Mammalian Protein Complex Structure, Function, and Membership Using Genome-Scale Fitness Screens. Cell Syst. 6, 555–568.e7. https://doi.org/10.1016/j.cels.2018.04.011.

Park, S.-Y., Schinkmann, K.A., and Avraham, S. (2006). RAFTK/Pyk2 mediates LPA-induced PC12 cell migration. Cell. Signal. 18, 1063–1071. https://doi.org/10.1016/j.cellsig.2005.08.018.

Perfetto, L., Briganti, L., Calderone, A., Perpetuini, A.C., Iannuccelli, M., Langone, F., Licata, L., Marinkovic, M., Mattioni, A., Pavlidou, T., et al. (2016). SIGNOR: A Database of Causal Relationships between Biological Entities. Nucleic Acids Res. 44, D548–D554. https://doi.org/10.1093/nar/gkv1048.

Rahman, M., Billmann, M., Costanzo, M., Aregger, M., Tong, A.H.Y., Chan, K., Ward, H.N., Brown, K.R., Andrews, B.J., Boone, C., et al. (2021). A method for benchmarking genetic screens reveals a predominant mitochondrial bias. Mol. Syst. Biol. 17. https://doi.org/10.15252/msb.202010013.

Rath, S., Sharma, R., Gupta, R., Ast, T., Chan, C., Durham, T.J., Goodman, R.P., Grabarek, Z., Haas, M.E., Hung, W.H.W., et al. (2021). MitoCarta3.0: an updated mitochondrial proteome now with sub-organelle localization and pathway annotations. Nucleic Acids Res. 49, D1541–D1547. https://doi.org/10.1093/nar/gkaa1011.

Rodchenkov, I., Babur, O., Luna, A., Aksoy, B.A., Wong, J.V., Fong, D., Franz, M., Siper, M.C., Cheung, M., Wrana, M., et al. (2020). Pathway Commons 2019 Update: integration, analysis and exploration of pathway data. Nucleic Acids Res. 48, D489–D497. https://doi.org/10.1093/nar/gkz946.

Saez-Rodriguez, J., Alexopoulos, L.G., Epperlein, J., Samaga, R., Lauffenburger, D.A., Klamt, S., and Sorger, P.K. (2009). Discrete logic modelling as a means to link protein signalling networks with functional analysis of mammalian signal transduction. Mol. Syst. Biol. 5, 331. https://doi.org/10.1038/msb.2009.87.

Sayan, B.S., Yang, A.L., Conforti, F., Tucci, P., Piro, M.C., Browne, G.J., Agostini, M., Bernardini, S., Knight, R.A., Mak, T.W., et al. (2010). Differential control of TAp73 and DeltaNp73 protein stability by the ring finger ubiquitin ligase PIR2. Proc. Natl. Acad. Sci. U. S. A. 107, 12877–12882. https://doi.org/10.1073/pnas.0911828107.

Schaefer, C.F., Anthony, K., Krupa, S., Buchoff, J., Day, M., Hannay, T., and Buetow, K.H. (2009). PID: The Pathway Interaction Database. Nucleic Acids Res. 37, D674–D679. https://doi.org/10.1093/nar/gkn653.

Sharp, R., Pyarelal, A., Gyori, B., Alcock, K., Laparra, E., Valenzuela-Escárcega, M.A., Nagesh, A., Yadav, V., Bachman, J., Tang, Z., et al. (2019). Eidos, INDRA, & Delphi: From Free Text to Executable Causal Models. In Proceedings of the 2019 Conference of the North American Chapter of the Association for Computational Linguistics (Demonstrations), (Minneapolis, Minnesota: Association for Computational Linguistics), pp. 42–47.

Shimada, K., Bachman, J.A., Muhlich, J.L., and Mitchison, T.J. (2021). shinyDepMap, a tool to identify targetable cancer genes and their functional connections from Cancer Dependency Map data. ELife 10. https://doi.org/10.7554/eLife.57116.

Sokolov, A., Carlin, D.E., Paull, E.O., Baertsch, R., and Stuart, J.M. (2016). Pathway-Based Genomics Prediction using Generalized Elastic Net. PLoS Comput. Biol. 12, e1004790. https://doi.org/10.1371/journal.pcbi.1004790.s012.

Steppi, A., Gyori, B., and Bachman, J. (2020). Adeft: Acromine-based Disambiguation of Entities from Text with applications to the biomedical literature. J. Open Source Softw. 5, 1708. https://doi.org/10.21105/joss.01708.

Surks, H.K., Richards, C.T., and Mendelsohn, M.E. (2003). Myosin phosphatase-Rho interacting protein. A new member of the myosin phosphatase complex that directly binds RhoA. J. Biol. Chem. 278, 51484–51493. https://doi.org/10.1074/jbc.M305622200.

Szklarczyk, D., Gable, A.L., Nastou, K.C., Lyon, D., Kirsch, R., Pyysalo, S., Doncheva, N.T., Legeay, M., Fang, T., Bork, P., et al. (2021). The STRING database in 2021: customizable protein–protein networks, and functional characterization of user-uploaded gene/measurement sets. Nucleic Acids Res. 49, D605–D612. https://doi.org/10.1093/nar/gkaa1074.

Torii, M., Arighi, C.N., Li, G., Wang, Q., Wu, C.H., and Vijay-Shanker, K. (2015). RLIMS-P 2.0: A Generalizable Rule-Based Information Extraction System for Literature Mining of Protein Phosphorylation Information. IEEE/ACM Trans. Comput. Biol. Bioinform. 12, 17–29. https://doi.org/10.1109/TCBB.2014.2372765.

Tsherniak, A., Vazquez, F., Montgomery, P.G., Weir, B.A., Kryukov, G., Cowley, G.S., Gill, S., Harrington, W.F., Pantel, S., Krill-Burger, J.M., et al. (2017). Defining a Cancer Dependency Map. Cell 170, 564–576.e16. https://doi.org/10.1016/j.cell.2017.06.010.

Tuncbag, N., Gosline, S.J.C., Kedaigle, A., Soltis, A.R., Gitter, A., and Fraenkel, E. (2016). Network-Based Interpretation of Diverse High-Throughput Datasets through the Omics Integrator Software Package. PLOS Comput. Biol. 12, e1004879. https://doi.org/10.1371/journal.pcbi.1004879.

Türei, D., Korcsmáros, T., and Saez-Rodriguez, J. (2016). OmniPath: guidelines and gateway for literature-curated signaling pathway resources. Nat. Methods 13, 966–967. https://doi.org/10.1038/nmeth.4077.

Valenzuela-Escárcega, M.A., Babur, Ö., Hahn-Powell, G., Bell, D., Hicks, T., Noriega-Atala, E., Wang, X., Surdeanu, M., Demir, E., and Morrison, C.T. (2018). Large-Scale Automated Machine Reading Discovers New Cancer-Driving Mechanisms. Database J. Biol. Databases Curation 2018. https://doi.org/10.1093/database/bay098.

Van Landeghem, S., Ginter, F., Van de Peer, Y., and Salakoski, T. (2011). EVEX: A PubMed-Scale Resource for Homology-Based Generalization of Text Mining Predictions. In Proceedings of BioNLP 2011 Workshop, (Portland, Oregon, USA: Association for Computational Linguistics), pp. 28–37.

Van Landeghem, S., Björne, J., Wei, C.-H., Hakala, K., Pyysalo, S., Ananiadou, S., Kao, H.-Y., Lu, Z., Salakoski, T., Van de Peer, Y., et al. (2013). Large-scale event extraction from literature with multi-level gene normalization. PloS One 8, e55814. https://doi.org/10.1371/journal.pone.0055814.

Vermehren-Schmaedick, A., Krueger, W., Jacob, T., Ramunno-Johnson, D., Balkowiec, A., Lidke, K.A., and Vu, T.Q. (2014). Heterogeneous Intracellular Trafficking Dynamics of Brain-Derived Neurotrophic Factor Complexes in the Neuronal Soma Revealed by Single Quantum Dot Tracking. PLoS ONE 9, e95113. https://doi.org/10.1371/journal.pone.0095113.

Wang, X., Ratnam, J., Zou, B., England, P.M., and Basbaum, A.I. (2009a). TrkB Signaling Is Required for Both the Induction and Maintenance of Tissue and Nerve Injury-Induced Persistent Pain. J. Neurosci. 29, 5508–5515. https://doi.org/10.1523/JNEUROSCI.4288-08.2009.

Wang, Y., Zheng, X.R., Riddick, N., Bryden, M., Baur, W., Zhang, X., and Surks, H.K. (2009b). ROCK isoform regulation of myosin phosphatase and contractility in vascular smooth muscle cells. Circ. Res. 104, 531–540. https://doi.org/10.1161/CIRCRESAHA.108.188524.

Wellbrock, C., Rana, S., Paterson, H., Pickersgill, H., Brummelkamp, T., and Marais, R. (2008). Oncogenic BRAF regulates melanoma proliferation through the lineage specific factor MITF. PloS One 3, e2734. https://doi.org/10.1371/journal.pone.0002734.

Westergaard, D., Stærfeldt, H.-H., Tønsberg, C., Jensen, L.J., and Brunak, S. (2018). A comprehensive and quantitative comparison of text-mining in 15 million full-text articles versus their corresponding abstracts. PLOS Comput. Biol. 14, e1005962. https://doi.org/10.1371/journal.pcbi.1005962.

Wick, K.R., Werner, E.D., Langlais, P., Ramos, F.J., Dong, L.Q., Shoelson, S.E., and Liu, F. (2003). Grb10 Inhibits Insulin-stimulated Insulin Receptor Substrate (IRS)-Phosphatidylinositol 3-Kinase/Akt Signaling Pathway by Disrupting the Association of IRS-1/IRS-2 with the Insulin Receptor *. J. Biol. Chem. 278, 8460– 8467. https://doi.org/10.1074/jbc.M208518200.

Wilcox, R.R. (1979). Estimating the Parameters of the Beta-Binomial Distribution. Educ. Psychol. Meas. 39, 527–535. https://doi.org/10.1177/001316447903900302.

Wong, J.V., Franz, M., Siper, M.C., Fong, D., Durupinar, F., Dallago, C., Luna, A., Giorgi, J., Rodchenkov, I., Babur, Ö., et al. (2021). Author-sourced capture of pathway knowledge in computable form using Biofactoid. ELife 10, e68292. https://doi.org/10.7554/eLife.68292.

Wooten, D.J., Gebru, M., Wang, H.-G., and Albert, R. (2021). Data-Driven Math Model of FLT3-ITD Acute Myeloid Leukemia Reveals Potential Therapeutic Targets. J. Pers. Med. 11, 193..

Wu, H., Zeinab, R.A., Flores, E.R., and Leng, R.P. (2011). Pirh2, a ubiquitin E3 ligase, inhibits p73 transcriptional activity by promoting its ubiquitination. Mol. Cancer Res. MCR 9, 1780–1790. https://doi.org/10.1158/1541-7786.MCR-11-0157.

Yuryev, A., Mulyukov, Z., Kotelnikova, E., Maslov, S., Egorov, S., Nikitin, A., Daraselia, N., and Mazo, I. (2006). Automatic Pathway Building in Biological Association Networks. BMC Bioinformatics 7, 171. https://doi.org/10.1186/1471-2105-7-171.

Zhang, H. (2004). The Optimality of Naive Bayes. In Proceedings of the Seventeenth International Florida Artificial Intelligence Research Society Conference (FLAIRS 2004), (Miami Beach, Florida, USA: AAAI Press), pp. 562–567.

Zucker, J., Paneri, K., Mohammad-Taheri, S., Bhargava, S., Kolambkar, P., Bakker, C., Teuton, J., Hoyt, C.T., Oxford, K., and Ness, R. (2021). Leveraging structured biological knowledge for counterfactual inference: A case study of viral pathogenesis. IEEE Trans. Big Data 7, 25–37..

